# Biogenesis of cytochromes *c* and *c*_1_ in the electron transport chain of malaria parasites

**DOI:** 10.1101/2024.02.01.575742

**Authors:** Aldo E. García-Guerrero, Rebecca G. Marvin, Amanda Mixon Blackwell, Paul A. Sigala

## Abstract

*Plasmodium* malaria parasites retain an essential mitochondrional electron transport chain (ETC) that is critical for growth within humans and mosquitoes and a key antimalarial drug target. ETC function requires cytochromes *c* and *c***_1_** that are unusual among heme proteins due to their covalent binding to heme via conserved CXXCH sequence motifs. Heme attachment to these proteins in most eukaryotes requires the mitochondrial enzyme holocytochrome *c* synthase (HCCS) that binds heme and the apo cytochrome to facilitate biogenesis of the mature cytochrome *c* or *c***_1_**. Although humans encode a single bifunctional HCCS that attaches heme to both proteins, *Plasmodium* parasites are like yeast and encode two separate HCCS homologs thought to be specific for heme attachment to cyt *c* (HCCS) or cyt *c***_1_** (HCC**_1_**S). To test the function and specificity of *P. falciparum* HCCS and HCC**_1_**S, we used CRISPR/Cas9 to tag both genes for conditional expression. HCC**_1_**S knockdown selectively impaired cyt *c***_1_** biogenesis and caused lethal ETC dysfunction that was not reversed by over-expression of HCCS. Knockdown of HCCS caused a more modest growth defect but strongly sensitized parasites to mitochondrial depolarization by proguanil, revealing key defects in ETC function. These results and prior heterologous studies in *E. coli* of cyt *c* hemylation by *P. falciparum* HCCS and HCC**_1_**S strongly suggest that both homologs are essential for mitochondrial ETC function and have distinct specificities for biogenesis of cyt *c* and *c***_1_**, respectively, in parasites. This study lays a foundation to develop novel strategies to selectively block ETC function in malaria parasites.

## INTRODUCTION

Malaria remains a devastating infectious disease, especially in sub-Saharan Africa ^1^. The lack of a long-lasting, broadly protective vaccine and increasing resistance to frontline antimalarial drugs underscore the urgency to unravel the basic biology of malaria parasites to identify new vulnerabilities suitable for therapeutic targeting. Severe malaria is caused by *Plasmodium falciparum* parasites, which infect and grow asexually within human red blood cells (RBCs) to cause all malaria symptoms (**Fig. 1**). RBCs contain abundant hemoglobin with approximately 1 x 10^9^ molecules of heme ^2^, and cellular metabolism involving this key cofactor is central to the biology of malaria parasites ^3–5^. During their 48-hour residence inside an RBC, parasites internalize and digest nearly 80% of the host-cell hemoglobin ^6^. This massive proteolysis releases copious free heme, which parasites detoxify by formation of hemozoin crystals inside their digestive vacuole. This critical pathway is a major antimalarial drug target and is central to the activity of chloroquine and artemisinin, two of the historically most successful antimalarial drugs ^7^.

**Figure 1.**
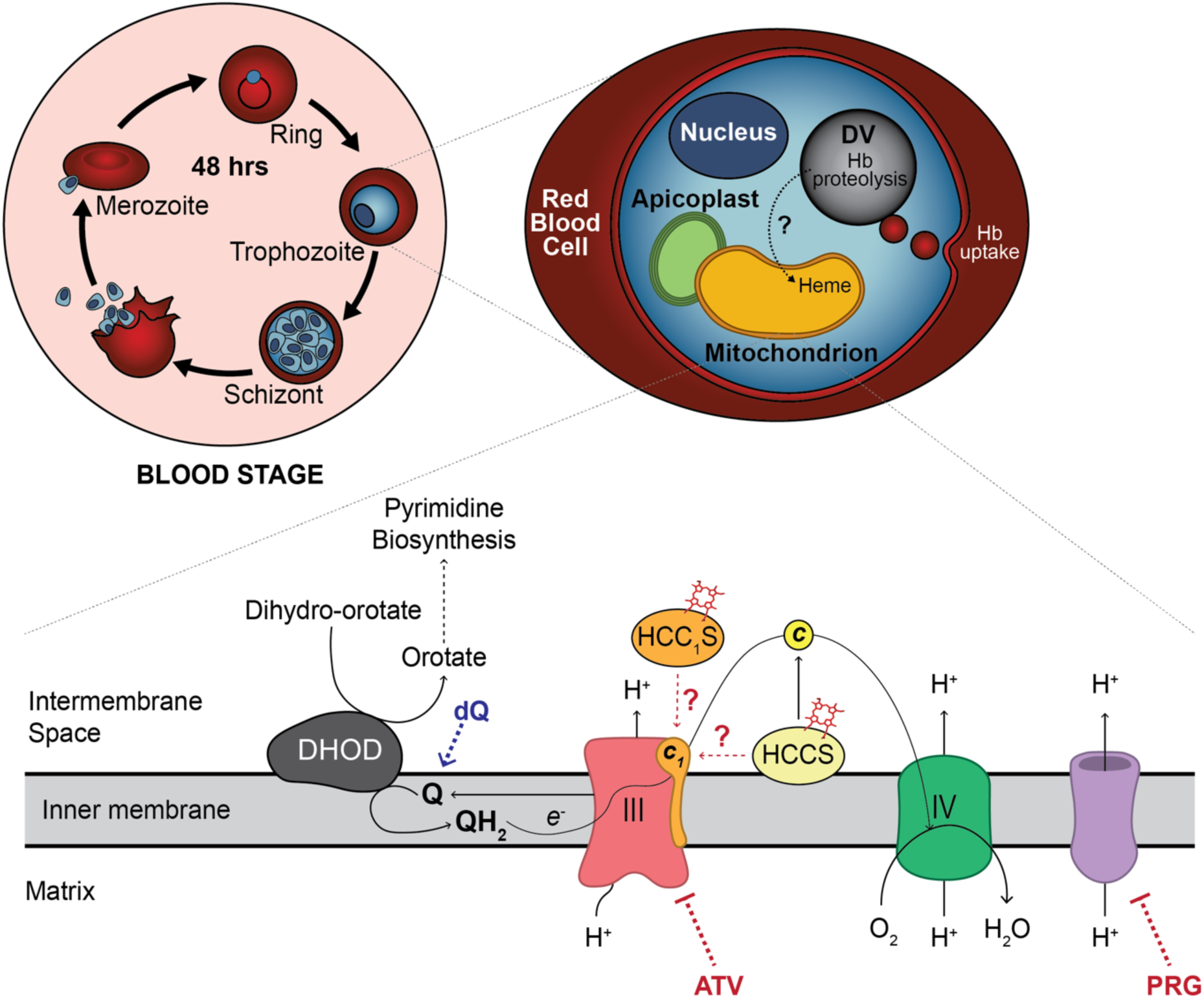
Schematic model of blood-stage infection and mitochondrial heme metabolism in *P. falciparum* parasites, focusing on the role of heme-dependent cytochromes *c* and *c*_1_ in the electron transport chain. ATV = atovaquone, DHOD = dihydroorotate dehydrogenase, dQ = decyl-ubiquinone, DV = digestive vacuole, Hb = hemoglobin, Q = oxidized ubiquinone, QH_2_ = reduced ubiquinol, III = Complex III/*bc*_1_, IV = Complex IV/Cyt *c* oxidase, *c* = cyt *c*, *c*_1_ = cyt *c*_1_, HCCS = holocytochrome *c* synthase, HCC_1_S, holocytochrome *c*_1_ synthase, PRG = proguanil. Purple = alternative, proguanil-sensitive pathway for maintaining transmembrane potential. Question marks indicate functional uncertainty.

*P. falciparum* parasites also require heme as a metabolic cofactor to support key aspects of their cellular physiology, especially the mitochondrial electron transport chain (ETC) that is essential for all stages of parasite development in humans and mosquitoes ^4, 5, 8^. ETC activity is directly inhibited by the current antimalarial drug, atovaquone ^9, 10^, and the key vulnerability of this pathway in all parasite stages has made it the subject of intense drug discovery efforts ^11–13^. Multiple heme-dependent cytochromes (cyt) are critical for ETC activity and parasite viability, including cyt *b* and *c***_1_** in Complex III, soluble cyt *c* that shuttles electrons between Complexes III and IV, and the heme A-binding Cox1 subunit of Complex IV ^4, 14^. In blood-stage parasites, the critical function of Complexes III and IV is oxidative recycling of the ubiquinone cofactor used as an electron acceptor by multiple mitochondrial dehydrogenases, especially dihydroorotate dehydrogenase required for pyrimidine biosynthesis (**Fig. 1**) ^10, 15–17^. Although electron transfer through these two complexes is accompanied by proton translocation across the inner membrane, this proton-pumping activity is not strictly essential for blood-stage parasites. *P. falciparum* has a redundant mechanism to polarize the mitochondrial inner membrane that is inhibited by proguanil, and blood-stage parasites rely on cytoplasmic glycolysis for ATP production rather than mitochondrial ATP synthase ^10, 18^. Of the known mitochondrial cytochromes, cyt *b* and Cox1 are encoded on the mitochondrial genome, which is not currently accessible to genetic modification. Cyt *c* (Pf3D7_1404100) and *c***_1_** (Pf3D7_1462700), however, are encoded in the nucleus, and we recently demonstrated that both cytochromes are critical for blood-stage parasites and ETC function ^17^. *P. falciparum* and other apicomplexan parasites also encode a divergent cyt *c* paralog (cyt *c*-2, Pf3D7_1311700) that binds heme but whose function is dispensable for blood-stage parasites ^17^.

Mitochondrial cyt *c* and *c***_1_**, which are retained by nearly all aerobic eukaryotes, are distinguished from most heme proteins by their use of a conserved CXXCH sequence motif to covalently bind heme via formation of stable thioether bonds between the Cys residues and the heme vinyl groups ^19–22^. In most eukaryotes, the assembly of these heme-bound cytochromes requires the enzyme holocytochrome *c* synthase (HCCS) that is located on the outer leaflet of the inner mitochondrial membrane and whose activity has been best studied in yeast and humans ^23,24^. Although hemylation of cyt *c* and *c***_1_** is central to eukaryotic respiration, no structure has been determined experimentally for any mitochondrial HCCS from any organism. Nevertheless, recombinant expression and functional reconstitution of cyt *c* hemylation by human HCCS have enabled a proposed mechanism for heme attachment ^24, 25^. According to this mechanism, HCCS binds heme via a conserved His residue, binds the N-terminal α-helix of apo cyt *c*, and positions the bound heme proximal to the CXXCH residues to facilitate stereospecific heme attachment and thioether bond formation to cyt *c.* These steps are followed by folding and release of mature cyt *c*. Although humans and other metazoans retain a single bifunctional HCCS that can hemylate both cyt *c* and *c***_1_** ^26^, other eukaryotes express two distinct HCCS homologs that preferentially hemylate cyt *c* (HCCS) or cyt *c***_1_** (HCC**_1_**S) ^23, 27^. Nevertheless, studies in *Saccharomyces cerevisiae* ^26^ and the apicomplexan parasite *Toxoplasma gondii* ^28^ have suggested that HCCS in these organisms has limited promiscuous activity in hemylating cyt *c***_1_**, in addition to its preferred cyt *c* substrate.

*P. falciparum* also expresses both HCCS (Pf3D7_1224600) and HCC**_1_**S (Pf3D7_1203600) homologs, but their functional properties have been sparsely studied. Genome-wide and targeted knock-out studies in *P. falciparum* and *P. berghei* have reported that both genes are refractory to disruption ^29–31^, suggesting non-redundant essential functions. Our prior work localized both proteins to the mitochondrion and also indicated that parasite HCCS but not HCC**_1_**S could hemylate cyt *c* or *c*-2 when expressed heterologously in *E. coli*, suggesting that each homolog is specific for hemylating cyt *c* or *c***_1_**, respectively ^17^. To directly test the function and specificity of HCCS and HCC**_1_**S in *P. falciparum* parasites, we used CRISPR/Cas9 to tag each gene for conditional knockdown studies. Loss of HCC**_1_**S impaired cyt *c***_1_** biogenesis and resulted in lethal ETC dysfunctions that were not rescued by HCCS over-expression. Knockdown of HCCS led to a more modest phenotype but strongly sensitized parasites to mitochondrial depolarization by proguanil, unmasking a key ETC defect. These studies unravel non-overlapping critical functions for HCCS and HCC**_1_**S in biogenesis of cyt *c* and *c***_1_** in malaria parasites. Specificity differences between human and parasite HCCS homologs can provide a basis to develop selective inhibitors of ETC function in *Plasmodium*.

## RESULTS

### *P. falciparum* encodes HCCS and HCC_1_S proteins homologous to yeast and humans

To better understand the sequence features of the *Plasmodium* HCCS and HCC**_1_**S homologs, we aligned these two proteins with HCCS orthologs from humans, yeast, and *Toxoplasma gondii* (**Fig. 2A**) and observed sequence identities ranging from 30-45%. *P. falciparum* HCCS (Pf3D7_1224600) had similar 35% and 38% sequence identity to yeast HCCS and HCC**_1_**S, respectively. The parasite HCC**_1_**S (Pf3D7_1203600), however, had higher sequence identity with yeast HCC**_1_**S (41%) than yeast HCCS (30%), consistent with its predicted specificity for cyt *c***_1_** and lack of detectable hemylation of cyt *c* in prior heterologous studies in *E. coli* ^17^. We also generated a maximum likelihood phylogenetic tree, based on alignment of the *Plasmodium* HCCS and HCC**_1_**S proteins with 52 orthologs from diverse eukaryotes, that further supported functional divergence between the parasite homologs (**Fig. 2B**).

**Figure 2.**
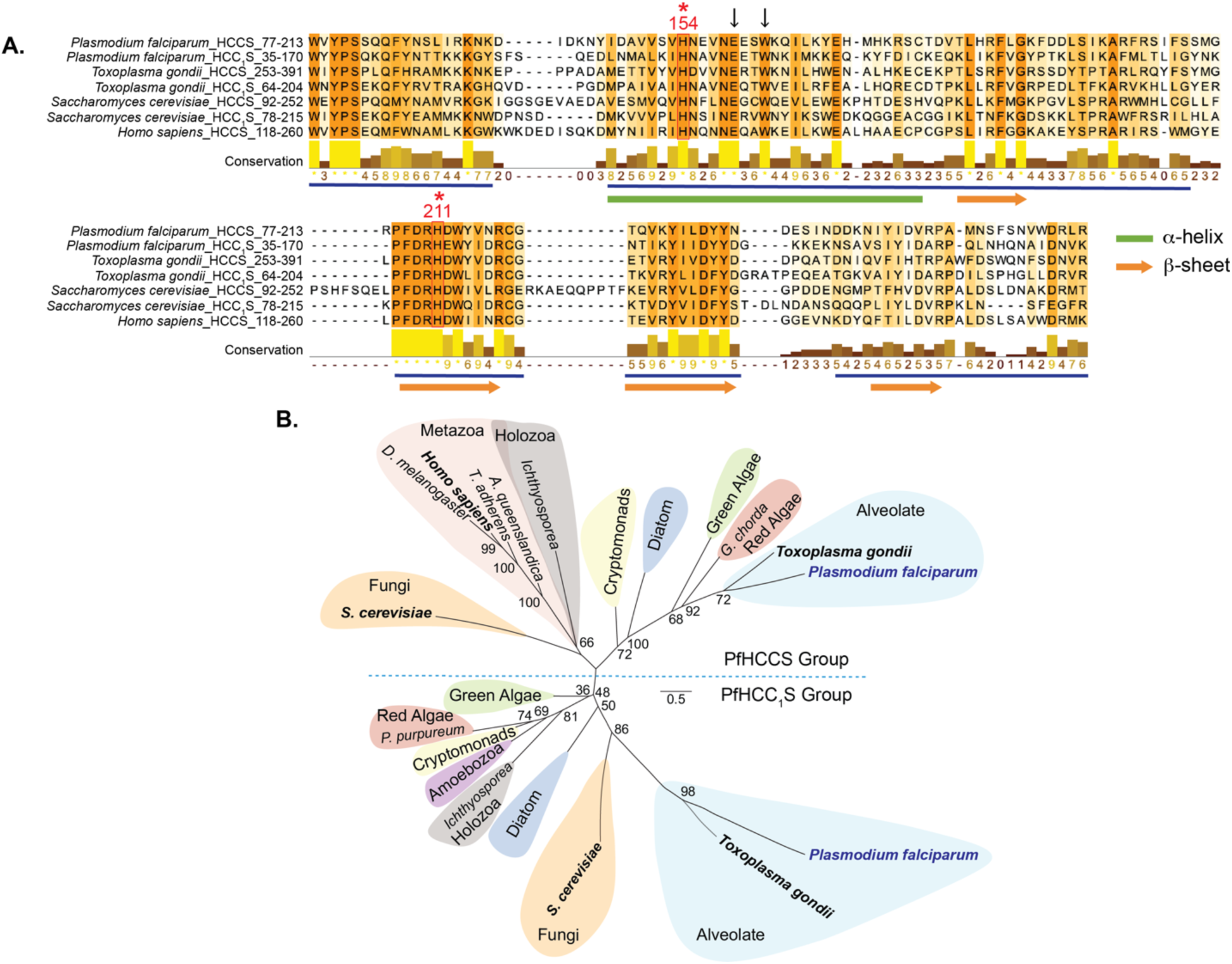
Sequence analysis of HCCS homologs. (A) CLUSTAL Sequence alignment of *P. falciparum* HCCS (77-213) and HCC_1_S (35-170) with homologs from *T. gondii* (HCCS: TGME49_293390, 92-252, and HCC_1_S: TGME49_314042, 64-204), *S. cerevisiae* (HCCS: CYC3/YAL039C, 92-252; and HCC_1_S: CYT2/YKL087C, 78-215), and *H. sapiens* (NP_001116080.1, 118-260). Conservation was measured by JalView with high scores (>5) underlined in blue. Vertical arrows and red asterisks highlight key conserved residues. For simplicity, N-terminal leader sequences are not shown. Residues included in alignments are indicated. (B) Molecular phylogeny of *P. falciparum* HCCS and HCC_1_S. BLAST analysis was used to identify orthologs of *P. falciparum* HCCS and HCC_1_S, with alignment of 52 sequences (Table S1) for construction of a Maximum Likelihood phylogenetic tree using the IQ-TREE and FigTree software. Bootstrap values are indicated at branch points. The scale bar corresponds to the expected sequence changes per 100 residues.

Based on the alignment in Fig. 2A and in agreement with prior studies ^24, 32^, we observed multiple areas of strong sequence conservation between all orthologs that included the conserved His residues at H154 and H211 (Human HCCS numbering) and neighboring residues E159 and W162. H154 has previously been demonstrated in human HCCS to serve as a key axial ligand required for heme binding by HCCS and hemylation of cyt *c*, but a functional role for H211 remains undefined ^24^. E159 and W162 also contribute to heme binding, and E159 mutations underpin the human disease called micropthalmia with linear skin defects syndrome ^32, 33^. The high conservation in these and nearby residues between human and *P. falciparum* HCCS is consistent with our prior observation that each HCCS ortholog could promiscuously catalyze heme attachment to cyt *c* from both organisms ^17^.

No three-dimensional structure has been determined experimentally for a eukaryotic HCCS enzyme. As an initial step towards understanding structural features of HCCS, the positioning of conserved residues, and the possible orientation of HCCS bound to cyt *c* and its CXXCH heme-binding motif, we first turned to available AlphaFold models for human HCCS and *P. falciparum* HCCS and HCC**_1_**S. Superposition of these structural models indicated very similar predicted structures for all three orthologs (**Fig. S1**), consistent with their substantial sequence similarity. The AlphaFold models suggest a central β-sheet that packs against an extended α-helix to create a central hydrophobic pocket, with several shorter α-helices predicted to pack around this hydrophobic core (**Fig. 3A and Fig. S1**). This predicted structure differs from a recent structural model for human HCCS generated by Rosetta and co-evolution approaches, which suggested a predominantly α-helical core structure ^25^. Nevertheless, both models predict that H154 is positioned within the protein core, where this residue is expected to coordinate bound heme ^24^. Indeed, AlphaFold 3 ^34^ modeled heme bound within the hydrophobic interior of parasite HCCS in an orientation coordinated by H154 (**Fig. S2**). The AlphaFold models further suggest that conserved H211 is positioned opposite H154 to possibly play a role as a second axial ligand for bound heme (**Fig. 3A and Fig. S2**). Prior mutagenesis studies with human HCCS found that H211 mutants showed minor reductions in heme attachment to cyt *c* that were less severe than H154 mutants ^24^, suggesting that H211 could contribute to heme binding to HCCS prior to association with apo cyt *c*. Such a role within the HCCS active site may underpin sequence conservation of H211 across phylogenetically disparate HCCS orthologs (**Fig. 2A**).

**Figure 3.**
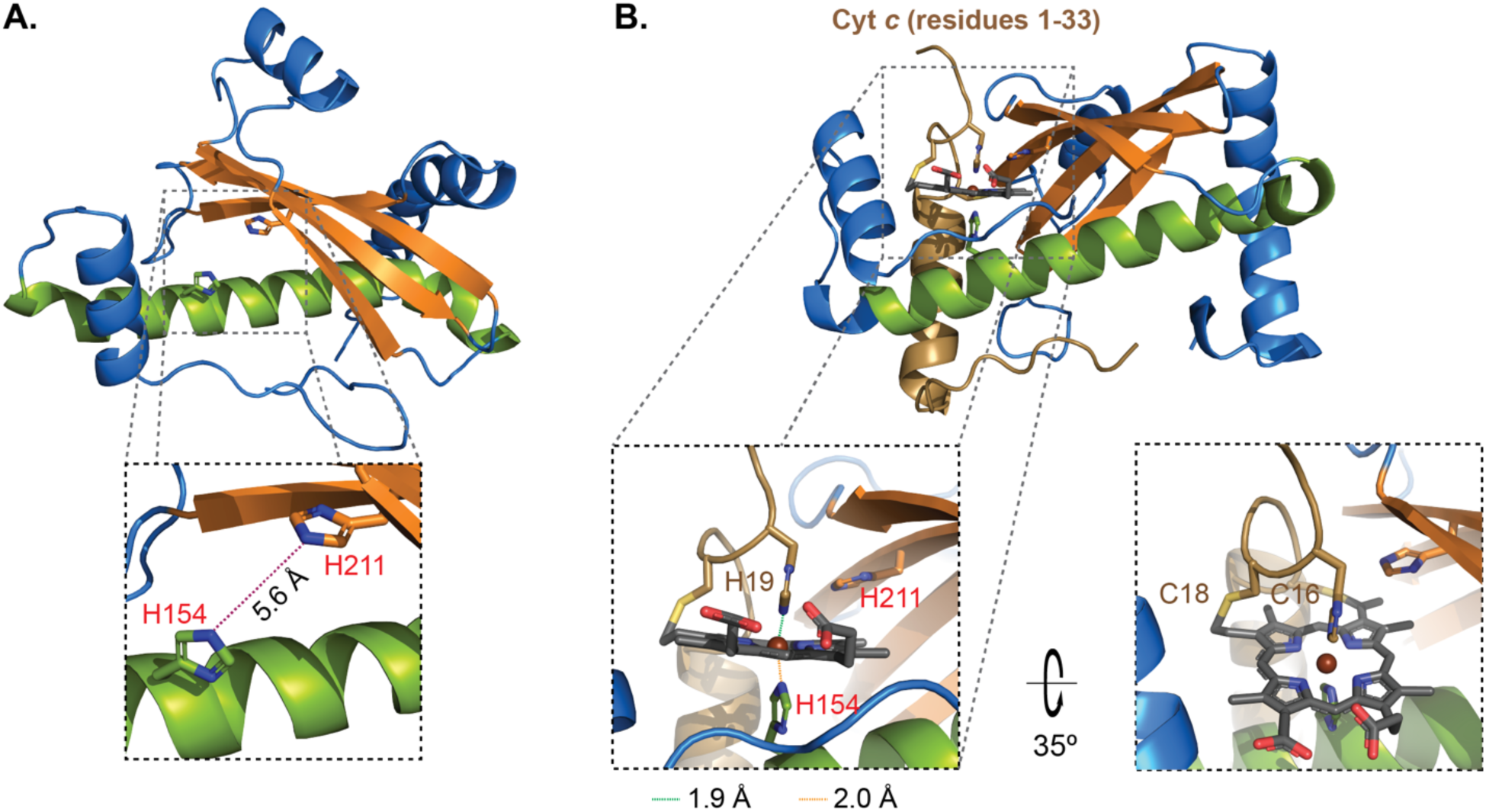
Structural modeling of HCCS. (A) AlphaFold model for PfHCCS showing proximity of conserved active-site His residues (H154, green; and H211, orange; human numbering). (B) AlphaFold 3 model of PfHCCS bound to heme and the N-terminal 33 amino acids of *P. falciparum* cyt *c* (Pf3D7_1404100) showing proximity of conserved His residues from HCCS (H154, green) and cyt *c* (H19, brown), using human sequence numbering.

We noted that AlphaFold 3 generated two distinct models for heme-bound HCCS, differing in the position of a central α-helix that either exposed or occluded the bound heme (**Fig. S2**). In both conformations, a conserved Lys residue (K133) was positioned within 2-4 Å of a heme propionate, suggesting a stabilizing electrostatic interaction that might underpin the strict conservation of this residue. The position of the central helix also appeared to pivot about a conserved Gly residue (G140), whose inherent conformational freedom may provide a flexible joint to enable rigid-body reorientation of this helix. The equilibrium distribution of these conformations gated by helical rearrangement may be critical for exposing the heme-bound HCCS active site for selective binding to the cyt *c* N-terminus during HCCS-catalyzed hemylation. Incisive tests of these hypotheses will require overcoming technical challenges to HCCS solubilization that have hampered experimental structure determination of this protein.

We next used AlphaFold 3 to model a possible complex between *P. falciparum* HCCS, heme, and the first 33 residues of the cyt *c* N-terminus, which include the N-terminal α-helix and CXXCH motif critical for recognition and heme attachment by HCCS ^24, 35^. In the modeled protein complex (**Fig. 3B**), which AlphaFold 3 generated based only on the input amino acid sequences and selecting heme as ligand, the N-terminal α-helix of cyt *c* packs against and perpendicular to one end of the central α-helix of HCCS. This orientation positions the conserved Cys residues of the CXXCH motif proximal to the active site cavity such that the conserved cyt *c* His residue (H19 in human numbering) extends into the HCCS active site and replaces H211 of HCCS in an orientation opposite H154. AlphaFold 3 modeled heme bound within the hydrophobic core of HCCS with the central iron coordinated by H154 of HCCS and the conserved H19 of cyt *c* such that the vinyl groups of heme were positioned proximal to C16 and C18 of cyt *c* (**Fig. 3B**). This model suggests a key intermediate in cyt *c* hemylation in which HCCS-bound heme is positioned by opposing His coordination to facilitate thioether bond formation between the Cys residues of cyt *c* and heme vinyl groups. This model is consistent with prior mechanistic suggestions for human HCCS maturation of cyt *c* based on biochemical experiments and structural modeling ^24, 25, 36^.

### HCC_1_S is essential for parasite viability, ETC function, and cyt *c*_1_ biogenesis

Prior genome-wide and targeted knockout studies in *P. falciparum* and *P. berghei* reported that both HCCS and HCC**_1_**S genes were refractory to disruption and thus likely essential ^29–31^, but neither protein has been directly studied in *Plasmodium falciparum*. We first focused on HCC**_1_**S, since our prior heterologous expression studies in *E. coli* suggested that this protein does not hemylate cyt *c* but were unable to specify a function for this protein in cyt *c***_1_** biogenesis ^17^. To directly test HCC**_1_**S function in *P. falciparum*, we used CRISPR/Cas9 to edit the HCC**_1_**S gene in Dd2 parasites to encode a C-terminal HA-FLAG epitope tag fusion and the 10x aptamer/TetR-DOZI system for conditional protein expression ^37^. In this system, protein expression occurs in the presence of the non-toxic small molecule, anhydrotetracycline (aTc) but is repressed upon aTc washout. PCR analysis confirmed correct editing of the HCC_1_S gene in transfected parasites (**Fig. S3, A and B**).

To evaluate HCC_1_S expression and its down-regulation upon aTc washout, we performed western blot (WB) experiments to detect the HA-FLAG epitope tags. In repeated experiments that included mitochondrial isolations, we were unable to detect the endogenously tagged protein, likely reflecting very low expression in blood-stage parasites and/or unusual solubility properties that resisted extraction in Triton X-100, SDS, and other detergents. To bypass this limitation, we used reverse transcription and quantitative PCR (RT-qPCR) to measure mRNA levels as a proxy for protein expression. The aptamer/TetR-DOZI system was designed to suppress translation by sequestering transcripts in P-bodies ^37^, and multiple prior studies have revealed a loss of transcript levels in –aTc conditions that suggests transcript degradation ^38, 39^. By RT-qPCR, we detected a strong reduction in transcript levels (∼70%) in -aTc compared to +aTc conditions that indicated robust down-regulation of HCC_1_S expression upon aTc washout (**Fig. 4A**). This downregulation in -aTc conditions was not observed for HCCS (which was not tagged for knockdown in this line), which instead displayed a modest increase in mRNA levels. To evaluate the impact of HCC_1_S knockdown on parasite growth, we synchronized the tagged parasites and evaluated their growth over 6 days (3 intraerythrocytic cycles) in ±aTc conditions. Parasites grew normally +aTc, but their growth was strongly suppressed after 3 days in -aTc conditions (**Fig. 4A**). By blood smear, we detected wide-spread evidence of parasite death at longer growth times (**Fig. S4A**), indicating that HCC_1_S function was essential for blood-stage growth.

**Figure 4.**
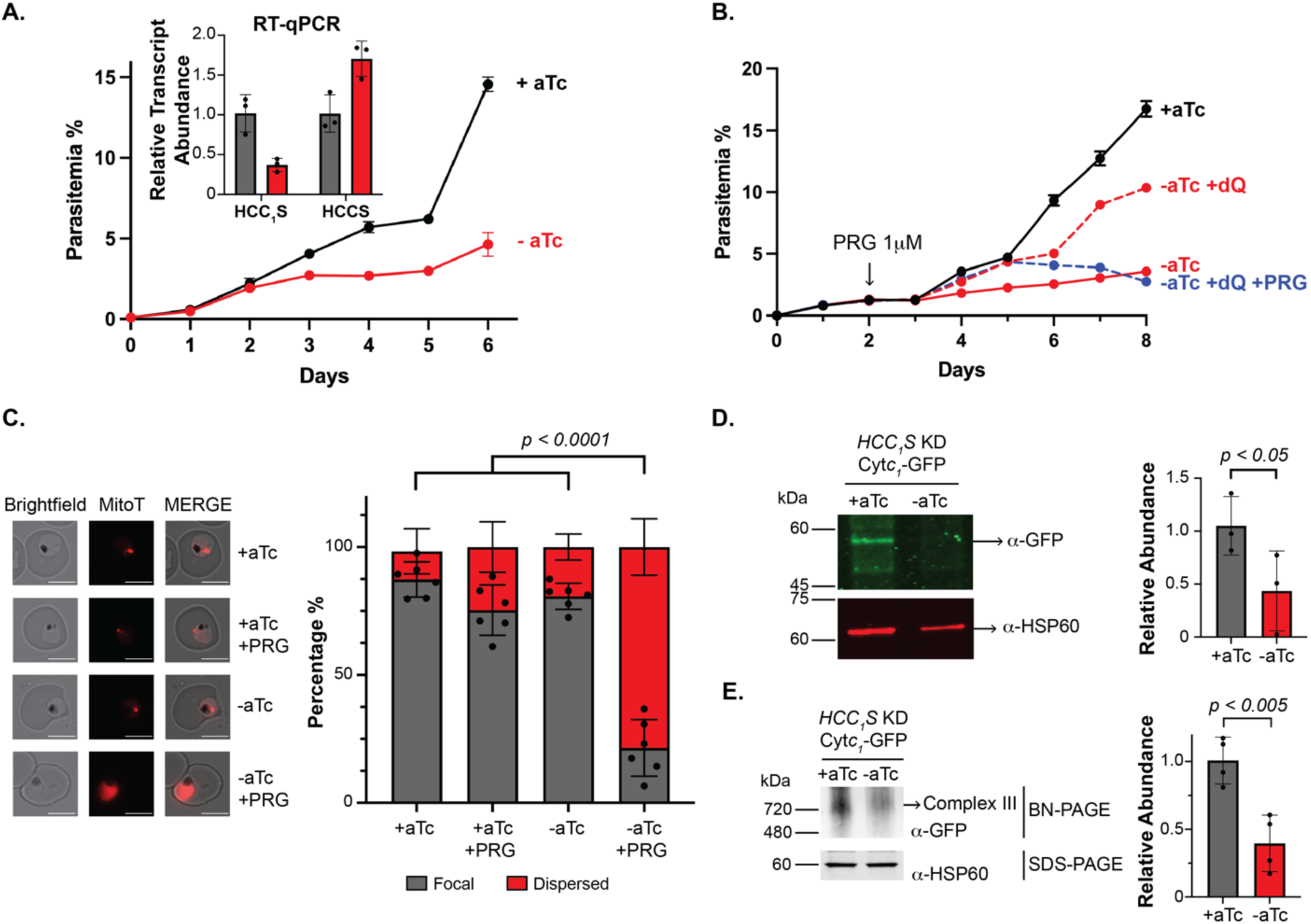
Functional impacts of HCC_1_S knockdown. (A) Synchronous growth assay of Dd2 parasites tagged with the aptamer/TetR-DOZI cassette with normal expression (+aTc) or inducible knockdown (-aTc) of HCC_1_S expression. Inset: RT-qPCR analysis of HCC_1_S and HCCS transcript levels (normalized to I5P, Pf3D7_0802500, and ADSL, Pf3D7_0206700) in biological triplicate samples cultured 4 days ±aTc in 15 µM decyl-ubiquinone (dQ). (B) Synchronous growth assay of HCC_1_S knockdown parasites cultured ±aTc, ±15 µM dQ, and ±1 µM proguanil (PRG, added day 2). Data points are the average ±SD of biological triplicates. Statistical analysis of growth differences is shown in Fig. S4C. (C) Fluorescence microscopy images of live HCC_1_S knockdown parasites cultured 5 days in 15 µM dQ, ±aTc, and ±1 µM proguanil and stained with 1 nM MitoTracker Red (MitoT). Scale bars represent 5 µm. Thirty parasites were imaged in each of 6 biological replicates to score MitoT staining as focal or dispersed and calculate the percentage of the total population in each condition with the indicated mitochondrial phenotype (average ±SD, *p* value from two-way ANOVA). (D) SDS-PAGE/WB analysis of cyt *c*_1_-GFP expression levels in HCC_1_S knockdown parasites cultured 3 days ±aTc/+dQ. Blot was probed with α-GFP and α-Hsp60 antibodies. Band intensities were quantified by densitometry, and the average normalized GFP:Hsp60 signal ±SD based on biological triplicate samples was plotted for ±aTc conditions (*p* value from two-tailed unpaired *t*-test). (E) Blue native-PAGE/WB analysis of cyt *c*_1_-GFP and SDS-PAGE/WB analysis of HSP60 expression levels in HCC_1_S knockdown parasites cultured 5 days ±aTc/+dQ. 20 µg of total protein from mitochondrial lysates were loaded for each lane. Blots were probed with α-GFP or α-HSP60 antibodies, with band intensities quantified by densitometry. The average normalized GFP intensity ±SD from biological replicates was plotted for ±aTc conditions. WB images from biological replicates and uncropped blots for (D) and (E) are shown in **Fig. S8** and **Fig. S9**, respectively.

Prior works have shown that blood-stage parasites can be rescued from defects in the mitochondrial electron transport chain by exogenous addition of the soluble ubiquinone analog, decyl-ubiquinone (dQ) ^15–17^. HCC_1_S knockdown is predicted to impair biogenesis of cyt *c*_1_, which is an integral and essential component of ETC Complex III. We thus predicted that dQ might rescue parasites from lethal loss of HCC_1_S (**Fig. 1**). To test this prediction, we supplemented parasite cultures with 15 µM dQ and monitored growth over 8 days ±aTc. We observed that dQ supplementation substantially rescued parasite growth in -aTc conditions (**Fig. 4B** and **Fig. S4C**), supporting a key role for HCC_1_S in ETC function. Addition of 1 µM proguanil (PRG), which inhibits the secondary pathway for mitochondrial polarization ^10, 18^, minimally impacted parasite growth in +aTc conditions (**Fig. S4B**) but negated dQ rescue and strongly impaired parasite growth in -aTc/+dQ conditions, as expected for defective ETC function upon HCC_1_S knockdown (**Fig. 4B**). Fluorescence microscopy of parasites grown for 5 days ±aTc/+dQ revealed that 1 µM proguanil specifically resulted in dispersed MitoTracker Red staining in -aTc conditions, which indicates mitochondrial depolarization (**Fig. 4C** and **Fig. S5**) ^10, 16, 17^. We conclude that HCC_1_S knockdown selectively impairs ETC function and sensitizes parasites to mitochondrial depolarization by proguanil, as expected for defective cyt *c*_1_ function.

Impairment of cyt *c*_1_ hemylation may reduce the folding stability and thus detectable levels of cyt *c*_1_ and/or its assembly into Complex III. To test the impact of HCC_1_S knockdown on cyt *c*_1_ levels, we tagged the HCC_1_S gene with the aptamer/TetR-DOZI system in Dd2 parasites that episomally expressed cyt *c*_1_-GFP ^16^ and confirmed integration by PCR (**Fig. S6, A and B**). Growth assays with synchronized parasites revealed a strong growth impairment in -aTc conditions (**Fig. S7**) that was very similar to the original HCC_1_S knockdown line (**Fig. 4A**). We harvested biological triplicate samples after 3 days of growth in ±aTc/+dQ conditions and fractionated samples by SDS-PAGE followed by western blot (WB) analysis of cyt *c*_1_-GFP, which migrates at ∼55 kDa due to N-terminal processing upon mitochondrial import ^16^ that is similar to prior observations for yeast and human cyt *c*_1_ ^40, 41^. This experiment revealed that HCC_1_S knockdown reduced cyt *c*_1_-GFP levels by ∼60% but did not impact mitochondrial Hsp60 levels (**Fig. 4D, Fig. S8,** and **Fig. S9**), supporting a key role for HCC_1_S in cyt *c*_1_ maturation and stability. It is possible that the GFP tag may enhance the stability of apo cyt *c*_1_ and thus mask a stronger reduction in cyt *c*_1_ levels upon HCC_1_S knockdown that would be observed in the absence of the GFP tag. Indeed, N- or C-terminal tags are commonly used to enhance the solubility and/or stability of fusion proteins during recombinant expression ^42, 43^.

To test the impact of HCC_1_S knockdown on cyt *c*_1_ incorporation into Complex III, we harvested samples after 5 days of growth in ±aTc/+dQ conditions and fractionated them by blue-native PAGE followed by WB analysis. Anti-GFP staining revealed a strong band at ∼700 kDa in +aTc conditions (**Fig. 4E**, **Fig. S8,** and **Fig. S9**), which is the expected size for Complex III dimer and consistent with recent BN-PAGE detections of Complex III in *P. falciparum* and *Toxoplasma gondii* ^44–47^. HCC_1_S knockdown in -aTc conditions, however, reduced the intensity of the 700 kDa band by ≥65%, suggesting a substantial defect in cyt *c*_1_ assembly into Complex III. We conclude that HCC_1_S function is critical for cyt *c*_1_ biogenesis and assembly into ETC Complex III.

### Loss of HCCS sensitizes parasites to mitochondrial depolarization by proguanil

To directly test HCCS function in blood-stage *P. falciparum*, we used a similar CRISPR/Cas9 strategy to tag the HCCS gene in Dd2 parasites to encode a HA-FLAG fusion tag and the aptamer/TetR-DOZI system for conditional knockdown. We used genomic PCR to confirm successful integration and gene editing in clonal parasites (**Fig. S3, A and C**). As with HCC_1_S, we were unable to detect the HA-FLAG-tagged HCCS by WB analysis, suggesting similar low expression. However, RT-qPCR analysis of transcript levels after 3 days of synchronous growth ±aTc revealed a specific ∼70% reduction in HCCS mRNA levels (**Fig. 5A**) that was similar to that observed for down-regulation of HCC_1_S (**Fig. 4A**). Synchronous growth assays in ±aTc conditions revealed a modest growth defect in -aTc conditions in the third cycle that was less severe than observed for HCC_1_S but suggested possible defects in ETC function. To test if this more modest phenotype was specific to HCCS knockdown in Dd2, we also tagged HCCS for knockdown in NF54 parasites and observed a similar phenotype (**Fig. S10**).

**Figure 5.**
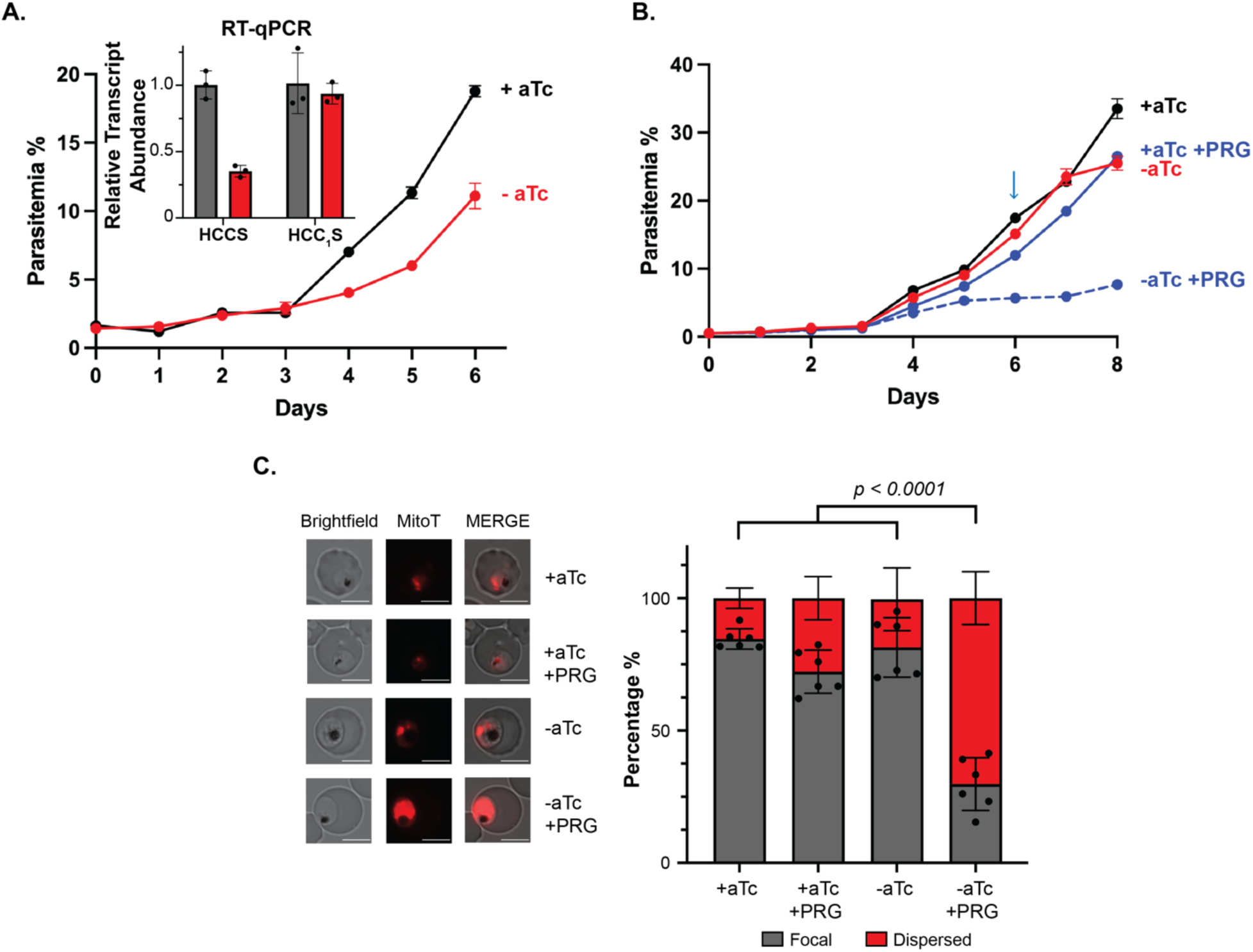
Functional impacts of HCCS knockdown. (A) Synchronous growth assay of Dd2 parasites (clone 1P10) tagged with the aptamer/TetR-DOZI cassette with normal expression (+aTc) or inducible knockdown (-aTc) of HCCS expression. Inset: RT-qPCR analysis of HCCS transcript levels (normalized to I5P and ADSL) in biological triplicates of synchronous parasites cultured 4 days ±aTc. (B) Synchronous growth assay of HCCS knockdown parasites cultured ±aTc and ±1 µM proguanil (PRG). All growth assay data points are the average ±SD of biological triplicates. Arrow indicates culture splits. (C) Fluorescence microscopy images of live HCCS knockdown parasites cultured 5 days ±aTc and ±1 µM proguanil and stained with 1 nM MitoTracker Red (MitoT). Thirty parasites were imaged in each of 6 biological replicates to score MitoT staining as focal or dispersed and calculate the percentage of the total population in each condition displaying the indicated mitochondrial phenotype (average ±SD). Significance was analyzed by two-way ANOVA to determine the indicated *p* values.

Addition of 1 µM proguanil had little or no impact on parasite growth +aTc but resulted in a stringent growth defect and parasite death in -aTc conditions (**Fig. 5B**). This observation suggests that HCCS knockdown results in a critical defect in proton translocation by Complex III and/or IV that is only unmasked in the presence of proguanil and inhibition of the secondary mechanism for mitochondrial polarization. Analysis by fluorescence microscopy, which revealed that proguanil selectively impaired mitochondrial accumulation of MitoTracker Red in -aTc conditions (**Fig. 5C** and **Fig. S5**), provided further support for this conclusion. Based on our prior observation that *P. falciparum* HCCS hemylates cyt *c* in heterologous expression studies in *E. coli* ^17^, we propose that HCCS knockdown in parasites impairs holo-cyt *c* biogenesis and reduces proton pumping by ETC Complex III and/or IV that sensitizes parasites to lethal mitochondrial depolarization by proguanil (**Fig. 1**). We further explore the phenotypic differences between HCC_1_S and HCCS knockdowns in the Discussion section below.

### *P. falciparum* HCCS and HCC_1_S have distinct functional specificities

Targeted gene knockout studies in *P. berghei* ^31^, prior biochemical experiments ^17^, and our observations in this study of parasite growth defects upon individual knockdown of HCC_1_S or HCCS (exacerbated by non-lethal proguanil) suggest essential and non-redundant functions and specificities for these proteins. To further test this conclusion and if over-expression of HCCS or HCC_1_S could rescue parasites from knockdown of either endogenous protein, we tagged the HCCS or HCC_1_S gene for conditional knockdown in the prior Dd2 parasite lines that episomally expressed HCCS-GFP or HCC_1_S-GFP ^17^ and confirmed correct integration by genomic PCR (**Fig. S6 and Fig. S11**). Synchronous growth assays revealed that parasites episomally expressing HCC_1_S-GFP were insensitive to knockdown of endogenous HCC_1_S and grew similarly in ±aTc conditions (**Fig. 6A**), as expected if episomal HCC_1_S-GFP functionally replaced the endogenous HCC_1_S. However, parasites episomally expressing HCCS-GFP displayed a severe growth defect upon HCC_1_S knockdown in -aTc conditions that was rescued by dQ (**Fig. 6B**), suggesting that HCCS is unable to functionally replace HCC_1_S. We observed the opposite pattern of rescue for loss of HCCS. Knockdown of endogenous HCCS resulted in no detectable growth impairment in parasites expressing HCCS-GFP (**Fig. 6C**) but a measurable growth defect in parasites expressing HCC_1_S-GFP (**Fig. 6D)** These observations support our conclusion that HCCS and HCC_1_S have distinct specificities for hemylation of cyt *c* and *c*_1_ that underpin non-redundant and critical functions.

**Figure 6.**
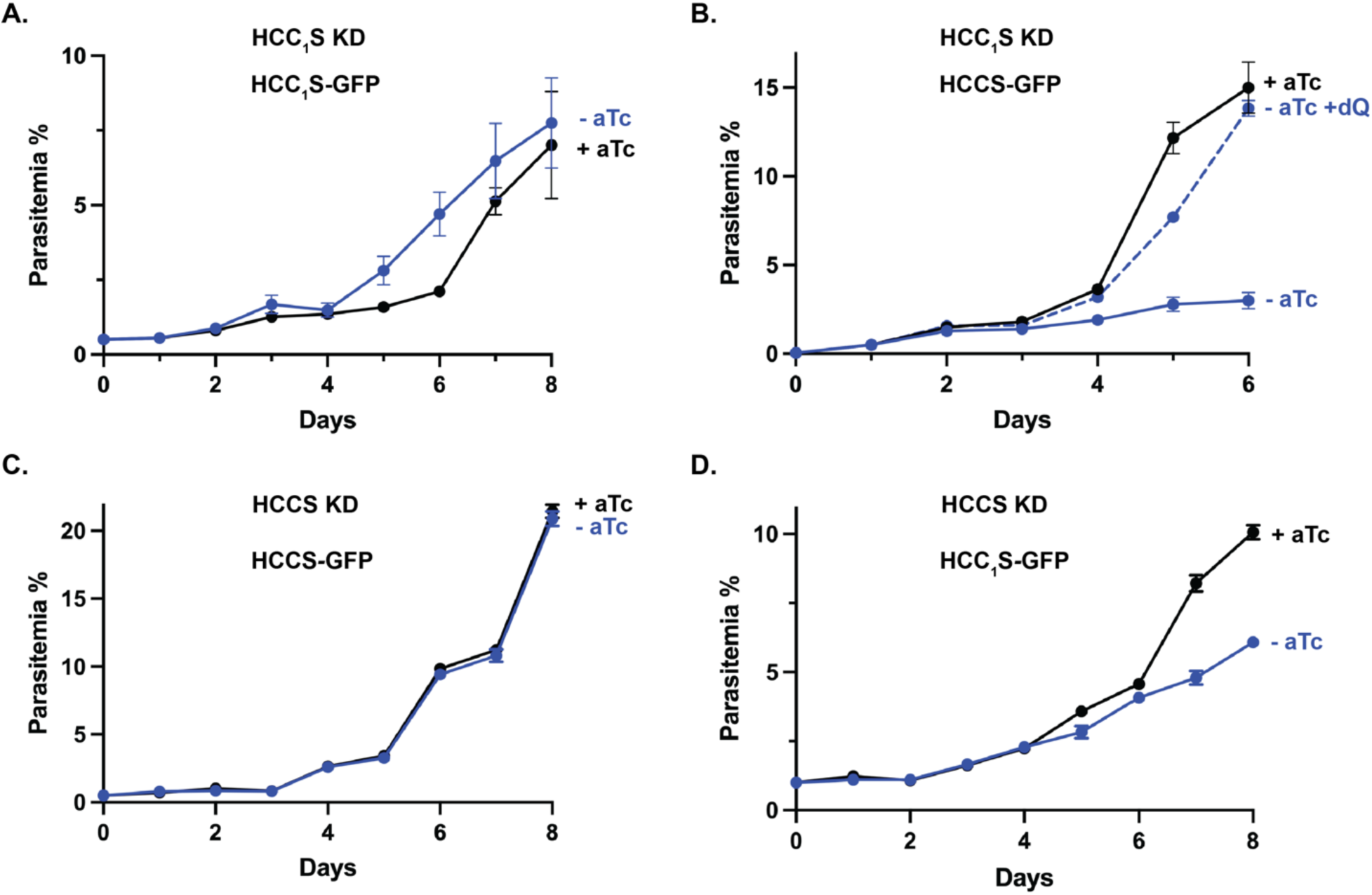
*P. falciparum* HCCS and HCC_1_S have non-redundant functions. Synchronous growth assays of HCC_1_S knockdown parasites episomally expressing HCC_1_S-GFP (A) or HCCS-GFP (B) or HCCS knockdown parasites episomally expressing HCCS-GFP (C) or HCC_1_S-GFP (D) and cultured in ±aTc and ±dQ (15 µM) conditions. All growth assay data points are the average ±SD of biological triplicate samples.

## DISCUSSION

The maturation and function of heme-bound cyt *c* and cyt *c*_1_ are central to mitochondrial electron transport chain activity in malaria parasites, especially respiratory Complexes III and IV (**Fig. 1**). Complex III is the molecular target of atovaquone ^9^ and has received considerable attention in the discovery and development of new antimalarial drugs ^11, 13^, especially as parasites that harbor atovaquone-resistant mutations are unable to complete mosquito-stage development and transmit to new hosts ^48, 49^. Prior work has also highlighted effective strategies to target parasites via Complex III during mosquito infection ^12^. These studies are nearly all based on targeting cyt *b* of Complex III, and few strategies have been identified to interfere with other essential ETC factors and functions. Recent efforts dissected the essential roles of cyt *c* and *c*_1_ for parasite ETC activity ^17^, but it is unclear how these protein functions might be directly targeted. In the present work, we identified critical independent functions for HCCS and HCC_1_S, required for covalent attachment of heme to cyt *c* and *c*_1_. This work lays a foundation for deeper structural and mechanistic studies to unravel differences within the heme-binding pockets of the human and parasite HCCS enzymes that can underpin selective therapeutic targeting of the *P. falciparum* ETC.

### Functional differences between *P. falciparum* HCCS and HCC_1_**S**

This study, in combination with prior functional tests based on heterologous reconstitution in *E. coli* ^17^, unveiled separate key functions for parasite HCCS and HCC_1_S in maturation of cyt *c* and *c*_1_, respectively (**Fig. 7**). Knockdown of HCC_1_S resulted in impaired cyt *c*_1_ biogenesis and incorporation into Complex III, lethal ETC dysfunction that stringently limited growth and could not be rescued by HCCS over-expression, and strong sensitivity to mitochondrial depolarization in the presence of proguanil. In comparison, HCCS knockdown caused a modest reduction in parasite growth that required combinatorial treatment with proguanil to stringently kill parasites via mitochondrial depolarization. It remains an important future challenge to define and understand the molecular differences between HCCS and HCC_1_S that confer specificity for hemylation of cyt *c* versus cyt *c*_1_. These functional differences may reflect sequence-specific features of each protein that underpin specific binding and/or heme attachment to cyt *c* or cyt *c*_1_. Alternatively, or in addition, there may be other cellular factors that interact with HCCS and HCC_1_S that contribute to substrate specificity.

**Figure 7.**
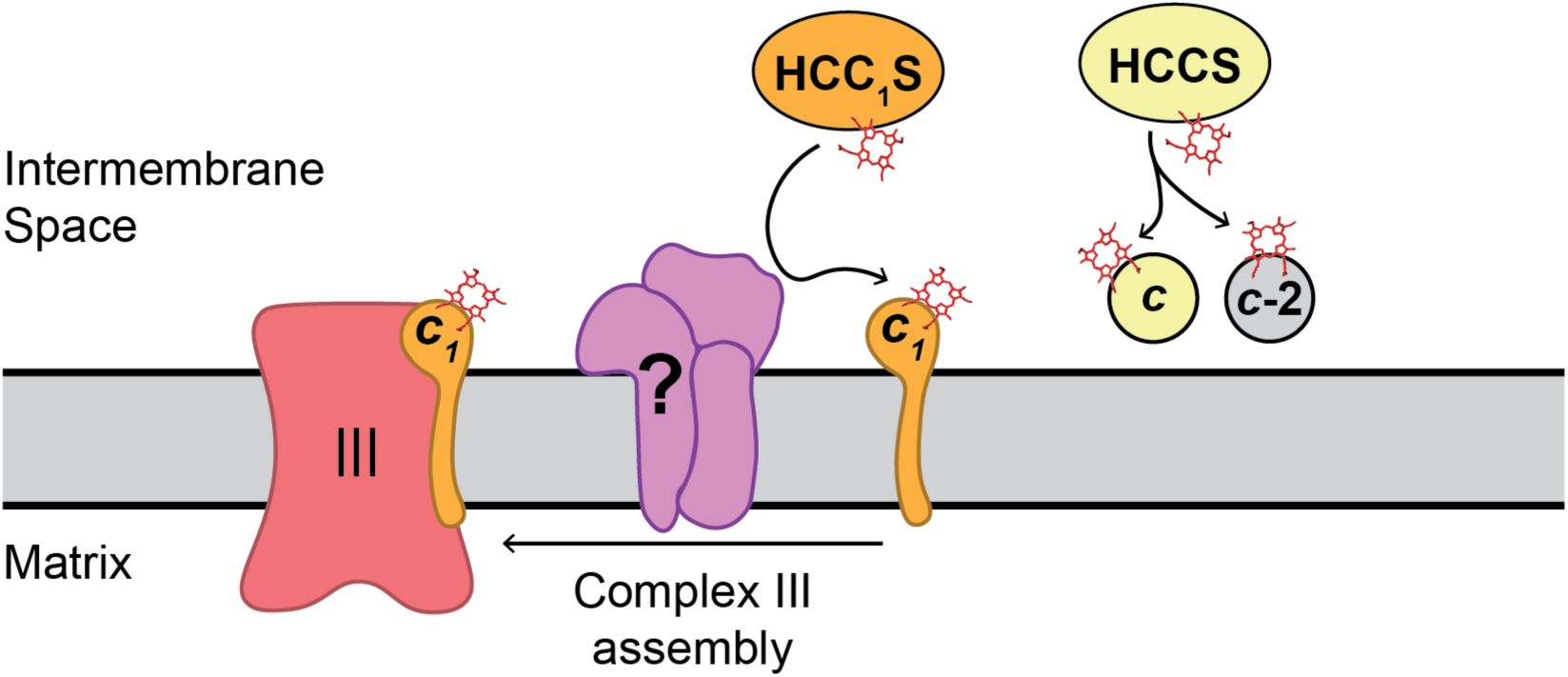
Schematic model for functional specificities of *P. falciparum* HCCS and HCC_1_S. The question mark indicates uncertainty in the mechanism of cyt *c*_1_ assembly into Complex III.

The strong growth defect due to loss of HCC_1_S phenocopies direct knockdown of cyt *c*_1_ ^17^ and underscores the tight dependence of ETC function and parasite viability on cyt *c*_1_ expression and heme-dependent maturation. This stringent coupling to cyt *c*_1_ may reflect both the direct function of cyt *c*_1_ in mediating critical electron transfer from the Rieske Fe-S protein to cyt *c* ^50^ and a secondary structural role played by cyt *c*_1_ in the assembly and integrity of Complex III ^51^, both of which are likely impaired by loss of HCC_1_S (**Fig. 7**). Indeed, our data provide direct evidence that HCC_1_S knockdown results in defective cyt *c*_1_ incorporation into Complex III (**Fig. 4E**). Prior work in yeast also highlighted that hemylation of cyt *c*_1_ by HCC_1_S is coupled to mitochondrial import and N-terminal processing ^40^. Import and processing of cyt *c*_1_ is sparsely studied in apicomplexan parasites. Nevertheless, recent work in *Toxoplasma* identified that the cyt *c*_1_ leader sequence that is cleaved upon mitochondrial import is incorporated into ATP synthase as a stable structural sub-unit ^45, 52^. It is not clear if this sub-unit is required for assembly and/or function of ATP synthase nor if this feature is conserved in *Plasmodium* parasites. Proteomic profiling of ATP synthase composition in *P. falciparum* did not report this cyt *c*_1_-derived subunit ^44^, although associations between cyt *c*_1_ and ATP synthase have been detected in blood-stage parasites ^53^. Nevertheless, this observation in *T. gondii* suggests an additional (and elegant) biochemical mechanism to couple cyt *c*_1_ import, hemylation, and maturation in apicomplexan parasites to broader assembly and function of the mitochondrial ETC and ATP synthase.

The more modest growth phenotype upon knockdown of HCCS compared to HCC_1_S, despite similar reductions in transcript abundance, is somewhat puzzling and may have multiple underpinnings. Cyt *c* function, which depends on heme attachment, is essential for ETC function and parasite viability ^17^. The mild phenotype for HCCS knockdown thus contrasts with the lethal phenotype for cyt *c* knockdown, the strong expectation of HCCS essentiality based on prior knockout studies ^29–31^, and the selective ability of HCCS (and not HCC_1_S) to hemylate cyt *c* in heterologous studies in *E. coli* ^17^. The first explanation we considered is that HCCS knockdown was insufficiently stringent such that residual expression was enough to support cyt *c* hemylation and attenuate the phenotypic severity of partial knockdown. However, HCCS knockdown sensitizes parasites to a much stronger, lethal phenotype when combined with 1 µM proguanil. This observation suggests a critical defect in proton pumping by Complexes III and/or IV due to HCCS knockdown, despite modest growth inhibition by itself, which can explain the synergistic lethality with proguanil which blocks the secondary pathway for mitochondrial polarization (**Figs. 1 and 7**).

Based on these observations, we favor an alternative model that HCCS knockdown in our current system impairs cyt *c* hemylation sufficiently to interfere with electron transport between Complexes III and IV but without substantially impairing oxidative recycling of ubiquinone by Complex III. A corollary of this model is that cyt *c*-mediated electron transport to Complex IV and concomitant proton pumping may be more tightly coupled to cyt *c* function than ubiquinone recycling by Complex III. This model predicts that a more stringent knockdown of HCCS, which might be achieved by adding a 5’ aptamer to the current 10x aptamers on the 3’ end of the HCCS gene, will impair ubiquinone recycling and result in a lethal growth phenotype that matches direct knockdown of cyt *c* and can be rescued by decyl-ubiquinone ^17^. We are currently testing this prediction.

### Implications for evolution of *P. falciparum* HCCS and HCC_1_S

Cyt *c* and *c***_1_** are retained by nearly all aerobic eukaryotes with a mitochondrion ^21, 22, 54^. Nature has evolved at least three distinct biochemical systems for *c*-type cytochrome hemylation, two of which are multi-protein systems that are unrelated evolutionarily to the single-protein HCCS system ^19, 20, 55^. HCCS and/or HCC**_1_**S proteins are broadly present in eukaryotes, including opisthokonts, alveolates, and amoebozoa ^55^. This broad retention across phylogenetically divergent organisms suggests that separate HCCS and HCC**_1_**S enzymes are ancient features of many eukaryotes. Within these clades that contain an HCCS homolog, metazoa and amoebozoa retain a single homolog (**Fig. 2B**) that appears to bifunctionally hemylate both cyt *c* and *c***_1_** ^20, 26^. For these organisms, low-level promiscuous hemylation of cyt *c***_1_** (in addition to the cognate cyt *c* substrate) was presumably enhanced in an ancestral organism such that a separate HCC**_1_**S enzyme became redundant and was lost ^20, 24^. Evidence for this model is provided by yeast, in which HCCS hemylates cyt *c* and less efficiently cyt *c***_1_** ^26^. As previously noted ^55^, the evolutionary origin of HCCS is unclear and remains a future challenge to understand. Indeed, our structural similarity searches of the RCSB Protein Data Bank with the AlphaFold-predicted *P. falciparum* HCCS structure using the Dali server ^56^ failed to identify substantial similarity to known protein structures.

Our results and those from prior biochemical ^17^ and gene-disruption studies ^29–31^ support a model in which both *P. falciparum* HCCS and HCC**_1_**S serve functionally independent and essential roles. This scenario contrasts with the related apicomplexan parasite *Toxoplasma gondii*, in which only the HCCS homolog appears to be essential for tachyzoite survival and shows evidence of hemylating both cyt *c* and *c*_1_, despite retention of an HCC_1_S paralog ^28^. Thus, the function, retention, and essentiality of HCCS and HCC_1_S homologs may be idiosyncratically tuned during evolution of different organisms depending on their unique cellular pressures for growth and survival. These functional differences may also involve additional cellular factors that influence observed specificities in vivo. Broader and deeper functional studies of these enzymes in differing organisms will facilitate understanding of the gain and loss of activities that underpin evolution of this enzymatic activity central to eukaryotic respiration.

### Source of heme for HCCS and HCC_1_S

*Plasmodium* parasites require heme as a metabolic cofactor for incorporation into mitochondrial cytochromes *b* and *c*_1_ in Complex III, cyt *c*, and cytochrome *a*/*a*_3_ in Complex IV ^3–5^. Parasites retain a complete biosynthesis pathway to make heme in the mitochondrion, but this pathway is dispensable for parasites during blood-stage growth and is only required during mosquito- and liver-stage growth ^57–62^. This observation implies that parasites can scavenge host-derived heme, although possible mechanisms for mobilizing labile heme released in the food vacuole during hemoglobin digestion and trafficking it to the mitochondrion remain undefined. HCCS and HCC_1_S function on the outer leaflet of the mitochondrial inner membrane ^63^ and must bind heme for subsequent attachment to cyt *c* and *c*_1_. Although proteomic studies in human cells suggest that the terminal heme synthesis enzyme, ferrochelatase, associates with other heme metabolism proteins at the mitochondrial inner membrane that may include or facilitate heme acquisition by HCCS ^64, 65^, parasite mechanisms for acquisition of biosynthetic or scavenged heme by HCCS and HCC_1_S are unknown. Our current study does not provide an answer to this question but lays a foundation for defining key interactions by these enzymes in *P. falciparum*, either through immunoprecipitation or proximity biotinylation coupled to tandem mass spectrometry, that can illuminate the heme-trafficking pathways that support biogenesis of heme-bound cyt *c* and *c*_1_.

## METHODS

### Cloning

Creation of pTEOE plasmids encoding HCCS-GFP and HCC_1_S-GFP was previously described ^17^. To tag HCCS and HCC_1_S for inducible knockdown, guide RNA sequences corresponding to TAACAAAAATTGTGAAAAAT and GCAAATGTGCAACAAACAAT (HCCS) and TTATTTATATAAAGATTTGT (HCC_1_S) were cloned into the pAIO CRISPR/Cas9 vector ^66^ using ligation-independent methods (primers HCCSgRNA1F, HCCSgRNA1R, HCCSgRNA2F, HCCSgRNA2R, HCC1SgRNA1F and HCC1SgRNA1R) by the Mutation Generation & Detection Core at the University of Utah. Donor repair plasmids for each gene were creating using assembly PCR to create a fused sequence containing a 3’ UTR homology region (HR) separated by an AflII site from a 5’ homology region that contained the 3’ end of the coding sequence (without stop codon) for each gene. This fused insert was then cloned into the pMG75 ^37^ plasmid using AscI/AatII cloning sites and ligation-independent methods and confirmed by Sanger sequencing. Donor plasmids to tag each gene were cloned from genomic DNA using the following primer sets: HCCS (3’ HR: primers 51/52, 5’ HR: primers 53/54) and HCC_1_S (3’ HR: primers 55/56, 5’ HR: primers 57/58). Primer sequences are given in Table S2.

### Parasite Culturing and Transfection

Experiments in this study were performed using *Plasmodium falciparum* Dd2 ^67^ and NF54 ^68^ parasites, whose identities were confirmed based on expected drug sensitivities and resistance. Parasites were *Mycoplasma*-free by PCR test. Dd2 parasites stably transfected with a pTEOE episome ^62^ encoding HCCS-GFP or HCC_1_S-GFP were available from a previously reported study ^17^ and cultured in 5 nM WR99210.

Parasites were cultured as previously described ^17, 39^, using Roswell Park Memorial Institute medium (RPMI-1640, Thermo Fisher 23400021) supplemented with 2.5 g/L AlbuMAX I Lipid-Rich BSA (Thermo Fisher 11020039), 15 mg/L hypoxanthine (Sigma H9636), 110 mg/L sodium pyruvate (Sigma P5280), 1.19 g/L HEPES (Sigma H4034), 2.52 g/L sodium bicarbonate (Sigma S5761), 2 g/L glucose (Sigma G7021), and 10 mg/L gentamicin (Invitrogen 15750060). Parasites were grown at 2% hematocrit in human red blood cells obtained from the University of Utah Hospital blood bank, at 37°C, and at 5% O_2_, 5% CO_2_, and 90% N_2_. Parasite-infected cultures were transfected in 1x cytomix ^69^ containing 50-100 µg midi-prep DNA by electroporation in 0.2 cm cuvettes using a Bio-Rad Gene Pulser Xcell system (0.31 kV, 925 µF). Transgenic parasites were selected based on plasmid resistance (blasticidin-S-deaminase) to 6 µM blasticidin-S (Gold Biotechnology B-800-100). Transgenic parasites modified with the aptamer/TetR-DOZI system or pTEOE episomes were cultured in 0.5-1 µM anhydrotetracycline (aTc, Cayman Chemicals 10009542) or 5 nM WR99210 (Jacobus Pharmaceuticals), respectively. Genetically modified parasites were genotyped by PCR. For growth assays with decyl-ubiquinone (dQ, Caymen Chemicals 55486005) and/or proguanil (Sigma 637321), the growth medium was supplemented with 15 µM dQ (≤0.3 % DMSO) and/or 1 µM proguanil (≤0.3 % DMSO).

For CRISPR/Cas9-based genome-editing experiments to tag HCCS (PF3D7_1224600) and HCC_1_S (PF3D7_1203600) to encode a C-terminal HA-FLAG epitope tag and the 10X aptamer/TetR-DOZI system for aTc-dependent expression ^37^, Dd2 or NF54 parasites were transfected in duplicate with 50 µg each of the linearized pMG74 donor plasmid (with gene-specific homology arms) and pAIO plasmid ^66^ (encoding Cas9 and gene-specific gRNA). Transfected parasites recovered for 48 hours in drug-free media and then were selected with 0.5 µM aTc and 6 µM blasticidin-S. Polyclonal parasites that returned from transfection were genotyped with primer sets 77_P1/54_P2, 53_P3/83_P4, 77_P1/PfHCCSRv7_P7 and 5’Hsp60Rv1_P8/PfHCCSRv9_P9 (HCCS) and 86_P1/58_P2, 57_P3/83_P4, 86_P1/PfHCC1SRv4_P7 and 5’Hsp60Rv1_P8/PfHCC1SRv4_P9 (HCC_1_S) to test for unmodified locus and successful integration, respectively. Polyclonal parasites that PCR analysis indicated were fully integrated without unmodified locus were used without cloning. Otherwise, clonal parasite integrants were obtained by limiting dilution. The Dd2 lines with HCC_1_S tagged for KD at its endogenous locus and episomally expressing cyt *c*_1_-GFP was created by first transfecting Dd2 parasites with cyt *c*_1_-GFP/pTEOE (selection with WR99210) and then transfecting with pAIO CRISPR/Cas9 and pMG74/donor repair plasmids to tag the HCC_1_S gene for knockdown (selection with blasticidin-S and aTc). Successful integration was confirmed by genotyping PCR as above. A similar strategy was used to tag HCCS or HCC_1_S for KD at its endogenous locus in DD2 parasites that episomally expressed HCCS-GFP or HCC_1_S-GFP and were previously reported ^17^. Primer sequences are given in Table S2.

### Parasite Growth Assays

For synchronous growth assays, asynchronous parasite cultures were treated with 5% D-sorbitol (Sigma S7900) for 10 minutes, resulting in a culture that contained ring-stage parasites. Sorbitol-treated cultures were then washed three times with RPMI media lacking aTc and divided into two equal parts before supplementing one part with 0.5 µM aTc. Parasites were diluted to 0.5% parasitemia ±aTc and allowed to grow over several days with daily media changes in biological triplicate samples. Parasitemia was monitored daily by flow cytometry by diluting 20 µL of each resuspended parasite culture well from each replicate into 200 µL of 1.0 µg/mL acridine orange (Invitrogen A3568) in phosphate buffered saline (PBS) pH 7.5 and analysis on a BD FACSCelesta system monitoring SSC-A, FSC-A, PE-A, FITC-A and PerCP-Cy5-5A channels. Parasitemia values are the average ± standard deviation of biological replicates. Growth assays in which parasitemia was plotted versus 2-day growth cycle was analyzed with an exponential growth model using GraphPad Prism 9.0. Observed differences in rate constants for exponential growth in different conditions were analyzed for significance by ANOVA using GraphPad Prism 9.0.

### RT-qPCR Analyses

Biological replicate cultures of HCCS or HCC_1_S knockdown Dd2 parasites were synchronized in 5% D sorbitol and grown for 72 hours in +aTc or -aTc/+dQ conditions prior to harvest. Four mL cultures at ∼10% parasitemia were harvested by centrifugation (2000 rpm for 3 min in a Beckman Coulter Allegra 25R table-top centrifuge) and stored at -20°C until use. Total RNA was isolated from frozen pellets and resuspended using a modified Trizol (Invitrogen) extraction protocol, as previously described ^39^. Washed and dried RNA pellets were resuspended in RNAse-free water, quantified by NanoDrop Spectrophotometer, and used immediately or stored at -80°C. One µg of RNA was DNAse-treated and reverse-transcribed using Superscript IV kit (Invitrogen) with the addition of 1 µM gene-specific reverse primers HCCSqR1, HCC_1_SqR1, GFPqR1, I5PqR1, and ADSLqR1. Subsequent cDNA was analyzed in technical triplicate through quantitative PCR using SYBR Green fluorescent probe with ROX (Invitrogen) in a QuantStudio 7 qPCR instrument (Applied Biosystems). C_p_ values for each target gene (primers HCCS_qF1, HCCS_qR1, HCC_1_S_qF1, and HCC_1_S_qR1) were normalized to average C_p_ values of the nuclear I5P gene (PF3D7_0802500, I5P_qF1 and I5P_qR1) and ADSL gene (Pf3D7_0206700, ADSL_qF1 and ADSL_qR1), then used to calculate relative RNA abundance values for each culture condition ±aTc, with normalization of the +aTc sample to a value of 1. Data is reported as the average ± SD of ≥3 biological replicates for each condition. All primer sequences are given in Table S2.

### Mitochondrial Isolation

Biological replicate 60 mL cultures of HCC_1_S KD Dd2 parasites were harvested at 15-18% parasitemia and treated with 0.05% saponin (Sigma 84510) in PBS (pH 7.2), incubated for 5 min at ambient temperature, and centrifuged at 3,000 x *g* for 30 min at 4°C. Isolated parasite-containing pellets were washed twice with PBS pH 7.2 and stored at -80°C until use. Parasite pellets were thawed on ice and resuspended with MEST buffer (20 mM Tris-HCl pH 7.4, 250 mM D-sorbitol, 1 mM EDTA pH 8.0, and EDTA-free protease inhibitor cocktail (Thermo Fisher J61852.XF). Parasites were lysed by 10 passages through a syringe fitted with a 25-gauge needle and 20 passages with a 27-gauge needle. The cellular debris and unlysed parasites were pelleted twice by cold centrifugation at 6,000 x *g* for 10 min at 4°C. The supernatant was centrifuged 15 min at 17,000 x *g* at 4°C to recover the crude mitochondrial fraction in the pellet. Total protein from the mitochondrial fraction was quantified using the Lowry method ^70^.

### SDS/Blue-Native PAGE and Western Blot Analyses

To evaluate expression levels of cyt *c*_1_-GFP in the HCC_1_S KD line ±aTc, biological replicate cultures of parasites were synchronized with 5% D-sorbitol to rings, grown for 72 hours +aTc or -aTc/+dQ, harvested as second-cycle trophozoites and schizonts, processed as described above to isolate mitochondria, and then analyzed by SDS-PAGE or blue native (BN)-PAGE.

For SDS-PAGE, 50 µg of mitochondrial protein was heated in 1X SDS sample buffer containing 5% fresh β-mercaptoethanol for 5 min at 65°C and loaded onto 12% polyacrylamide gels, fractionated by gel electrophoresis, and transferred to nitrocellulose membrane, as previously described ^16, 17^. Membranes were blocked at room temperature for 1 hour in 5% skim milk/PBS and probed overnight at 4°C with 1:1000 dilutions of rabbit anti-HSP60 (Novus NBP2-12734) and goat anti-GFP (Antibodies.com A121560) antibodies. Membranes were washed 3 times in TBS buffer supplemented with 0.1% TWEEN-20 (Millipore P1379), probed with 1:5,000 dilutions of donkey anti-rabbit IRDye680 (Licor 926-68023) and donkey anti-goat IRDye800CW (Licor 926-32214) 2° antibodies in TBST buffer, washed 3 times in TBST buffer, and imaged on a Licor Odyssey CLx imager. Band intensities were quantified by densitometry using ImageJ and analyzed by GraphPad Prism 9.0.

For BN-PAGE, 20 µg of mitochondrial protein was mixed with 1X Native-PAGE sample buffer (Invitrogen BN2003) supplemented with 1% digitonin and incubated on ice for 15 min. The sample was centrifuged at 17,000 x g for 10 min., and 20 µL of the supernatant was loaded and run on a NativePAGE 4-16% Bis-Tris gel (Invitrogen BN1002BOX) and transferred to PVDF membrane (Rio-Rad 1620177, pre-wetted in 70% methanol). The membrane was destained in 70% methanol and then washed with TBS-T, blocked, probed with goat anti-GFP 1° antibody, and washed as above. The membrane was then probed with a 1:5,000 dilution of rabbit anti-goat-HRP 2° antibody (Santa Cruz Biotechnology sc-2768), washed, developed using the SuperSignal West Pico PLUS enhanced chemiluminescence reagent (Thermo Fisher 34577), and imaged on an Invitrogen/Thermo Fisher iBright 1500 imager.

### Fluorescence Microscopy

For mitochondrial depolarization experiments with HCCS and HCC_1_S knockdown parasites, cultures were synchronized with 5% D-sorbitol, cultured +aTc or -aTc/+dQ for 120 hours before addition of 1 µM proguanil for 4 hours and 1 nM MitoTracker Red for 30 minutes, and imaged as third-cycle trophozoites/schizonts. 30 total parasites for each condition across 4 independent experiments were scored in a sample-blinded manner for focal or dispersed MitoTracker signal. Images were taken on DIC/brightfield, DAPI, and RFP channels using an EVOS M5000 imaging system. Images were processed using ImageJ with all brightness/contrast adjustments made on a linear scale.

### Alignment and Phylogenetic Analyses

The protein sequences for parasite HCCS (Pf3D7_1224600) and HCC_1_S (Pf3D7_1203600) were analyzed by PSI-BLAST ^71^ to identify orthologs in representative species of distinct organisms. Sequences (Table S1) were aligned by CLUSTAL Omega ^72^ and analyzed using Jalview ^73^. The multisequence alignment was uploaded to the IQ-TREE webserver (http://iqtree.cibiv.univie.ac.at) with ultrafast bootstrap analysis ^74^. The resulting maximum likelihood phylogenetic tree from 1000 bootstrap alignments was analyzed using FigTree (http://tree.bio.ed.ac.uk/software/figtree/). The raw phylogenetic tree is shown in Fig. S12.

### Structural Modeling

Structural models for human HCCS and *P. falciparum* HCCS and HCC_1_S were obtained from the AlphaFold Protein Structure Database (https://alphafold.ebi.ac.uk) ^75^. Structural models for parasite HCCS bound to heme without or with the N-terminal 33 residues of *P. falciparum* cyt c (PF3D7_1404100) were generated using AlphaFold 3 ^34^. Protein models were visualized and analyzed using PyMOL ^76^.

## SUPPORTING INFORMATION

AlphaFold models of HCCS (Figure S1), sequence conservation and conformational variation of HCCS (Figure S2), genome editing of HCC_1_S and HCCS in parasites (Figure S3), growth analysis of HCC_1_S knockdown parasites (Figure S4), additional live-parasite microscopy images of HCC_1_S and HCCS knockdown parasites cultured ±proguanil (Figure S5), genome editing of HCC_1_S in parasites transfected for episomal expression (Figure S6), growth assay of HCC_1_S knockdown parasites transfected for episomal expression of cyt c_1_-GFP (Figure S7), additional western blot images of HCC_1_S knockdown parasites episomally expressing cyt c_1_-GFP (Figure S8), uncropped western blot images (Figure S9), genotyping HCCS integration in NF54 parasites (Figure S10), genotyping HCCS integrations in parasites transfected for episomal expression (Figure S11), raw phylogenetic tree of HCCS and HCC_1_S homologs (Figure S12), list of organisms and HCCS or HCC_1_S protein sequences aligned in Figure S12 (Table S1), list of primers (Table S2) (PDF)

## ACKNOWLEDGMENTS

We thank Chris Hill, Robert Kranz, Tyler Starr, Henry Wienkers, Dennis Winge and members of the Sigala lab for helpful discussions. We thank Megan Okada for illustration assistance in making the scheme in Fig. 1. This work was supported by NIH grant R35GM133764 (PAS) and a Burroughs Wellcome Fund Career Award at the Scientific Interface (PAS). PAS is a Pew Biomedical Scholar, supported by The Pew Charitable Trusts. DNA synthesis and sequencing, fluorescence microscopy, creation of CRISPR/Cas9 reagents, and flow cytometry were performed using core facilities at the University of Utah and the Center for Iron and Heme Disorders.

---

**Figure S1.**
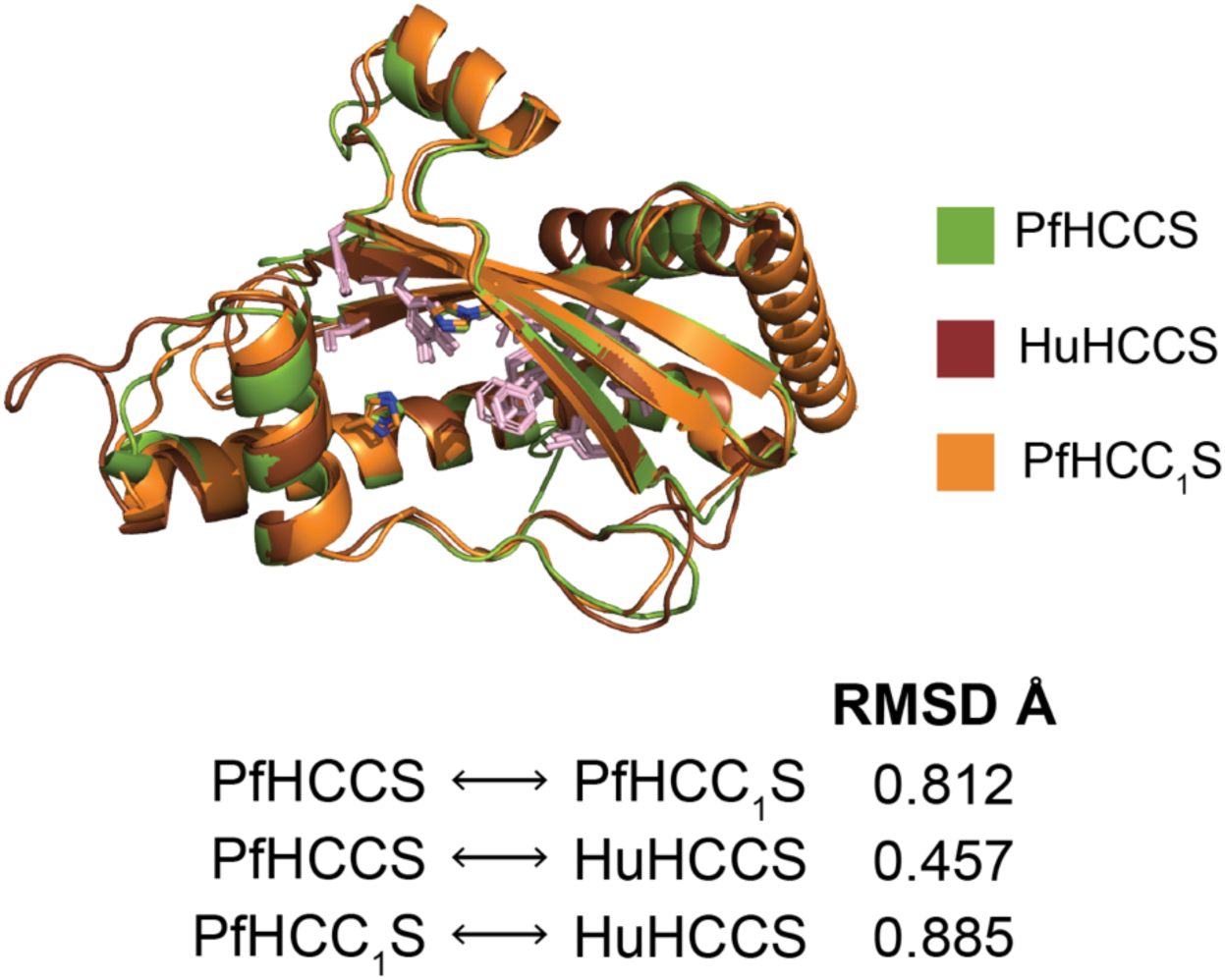
Structural alignment of the AlphaFold models *P. falciparum* HCCS, *P. falciparum* HCC_1_S, and human HCCS.

**Figure S2.**
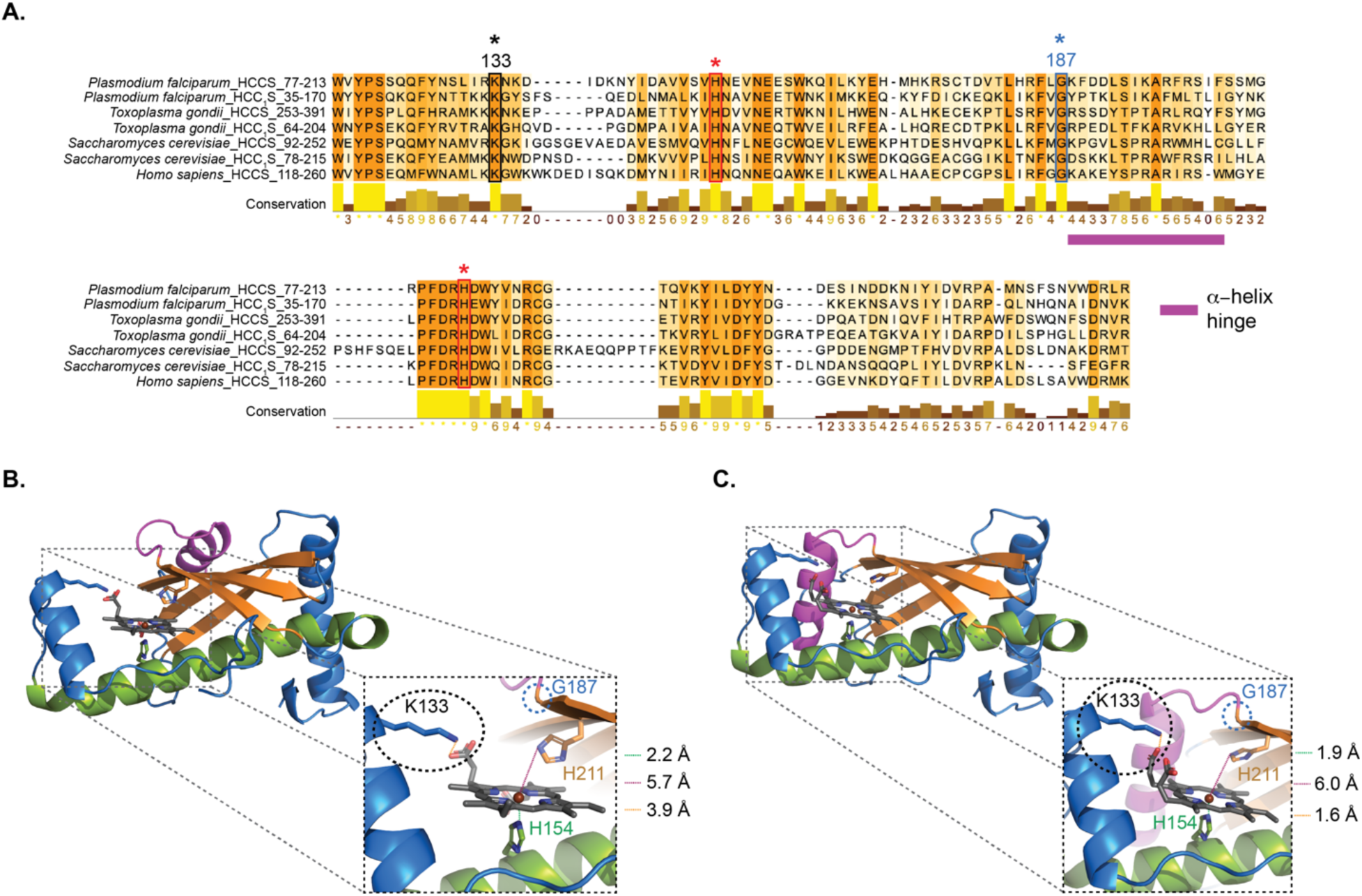
Sequence conservation and conformational variation of HCCS. (A) Sequence alignment of parasite, yeast, and human HCCS homologs highlighting conservation of K92 and G140 (human numbering). (B) and (C) AlphaFold-predicted structural models for *P. falciparum* HCCS with central alpha helix (purple) in two distinct conformations that expose (B) or occlude (C) the heme-binding pocket.

**Figure S3.**
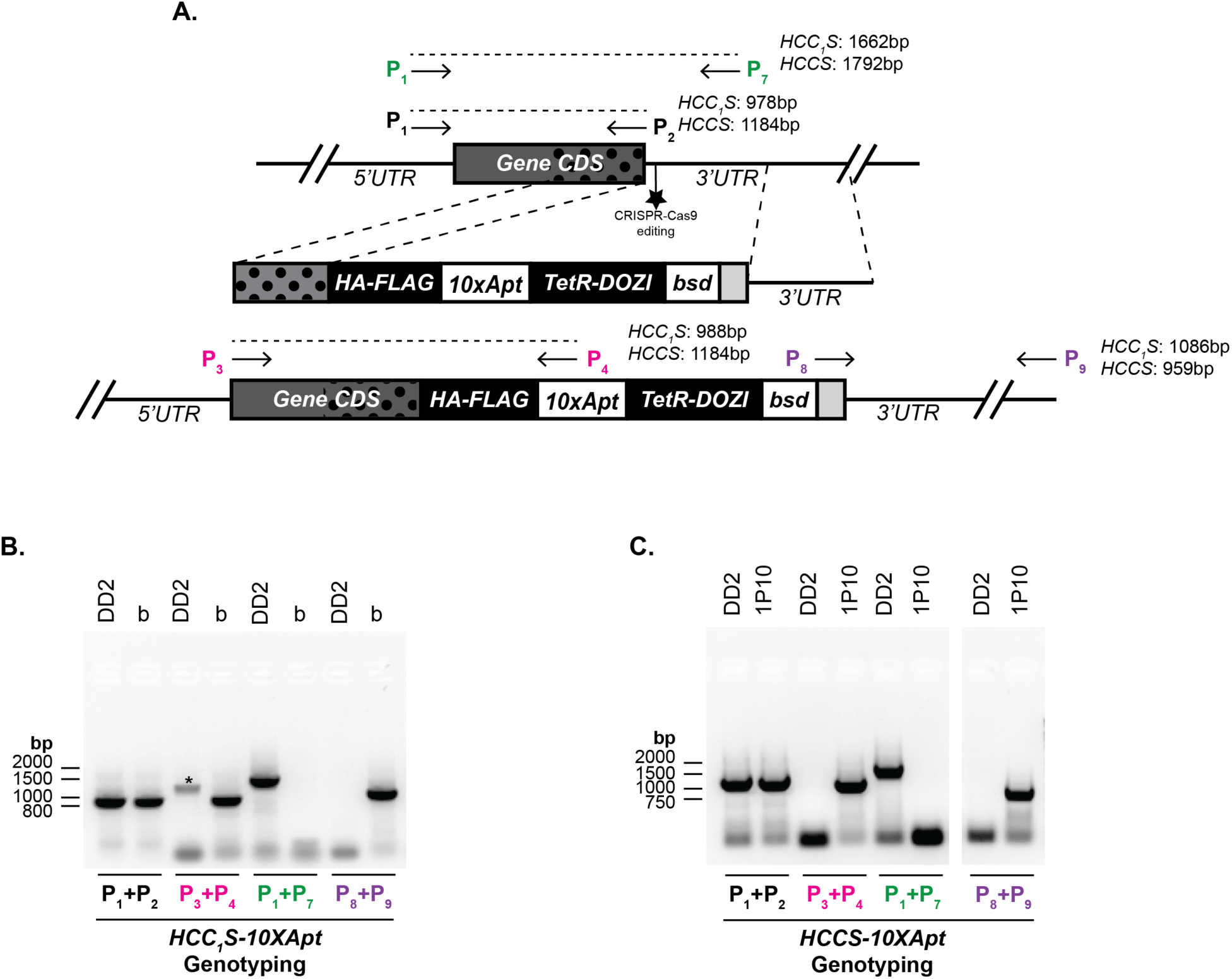
Genotyping HCC_1_S and HCCS integrations in Dd2 parasites. (A) Schematic model for editing the chromosomal gene for *P. falciparum* HCCS and HCC_1_S to encode a C-terminal HA-FLAG epitope tag fusion and the aptamer/TetR-DOZI system. PCR primer sites are indicated with a “P” along with expected amplicon sizes for primer pairs. (B) Genotype PCR for parental and polyclonal transfectants to edit HCC_1_S. Asterisk indicates non-specific amplicon. (C) Genotype PCR for parental and a clonal transfectant to edit HCCS.

**Figure S4.**
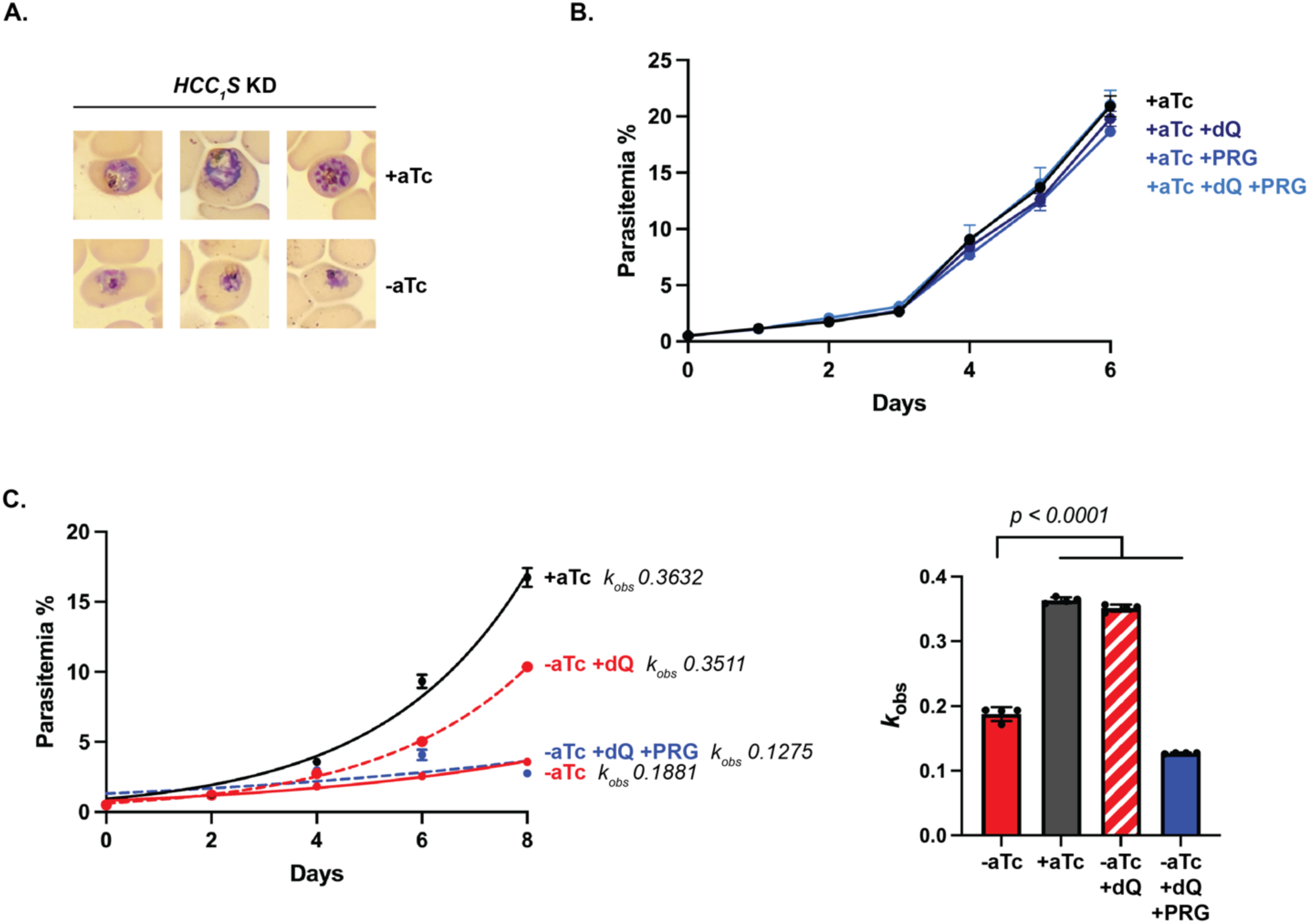
Growth analyses of HCC_1_S knockdown parasites. (A) Blood-smear analysis of synchronous HCC_1_S knockdown parasites grown 3 days in ±aTc conditions. (B) Synchronous growth analysis of HCC_1_S knockdown parasites cultured +aTc, ±15 µM decyl ubiquinone (dQ), and ±1 µM proguanil (PRG). (C) Synchronous growth assay of HCC_1_S knockdown parasites cultured ±aTc, ±15 µM dQ, and ±1 µM PRG (data are from Fig. 4B). Parasitemia values from even number days were fit to an exponential growth model in GraphPad Prism to determine the observed rate constant (*k*_obs_) for exponential growth. Growth rate constants were determined for each of the four biological replicate samples and used to determine the average ±SD rate constant for each growth condition. Statistical significance (with indicated p values) was determined by two-way ANOVA using GraphPad Prism.

**Figure S5.**
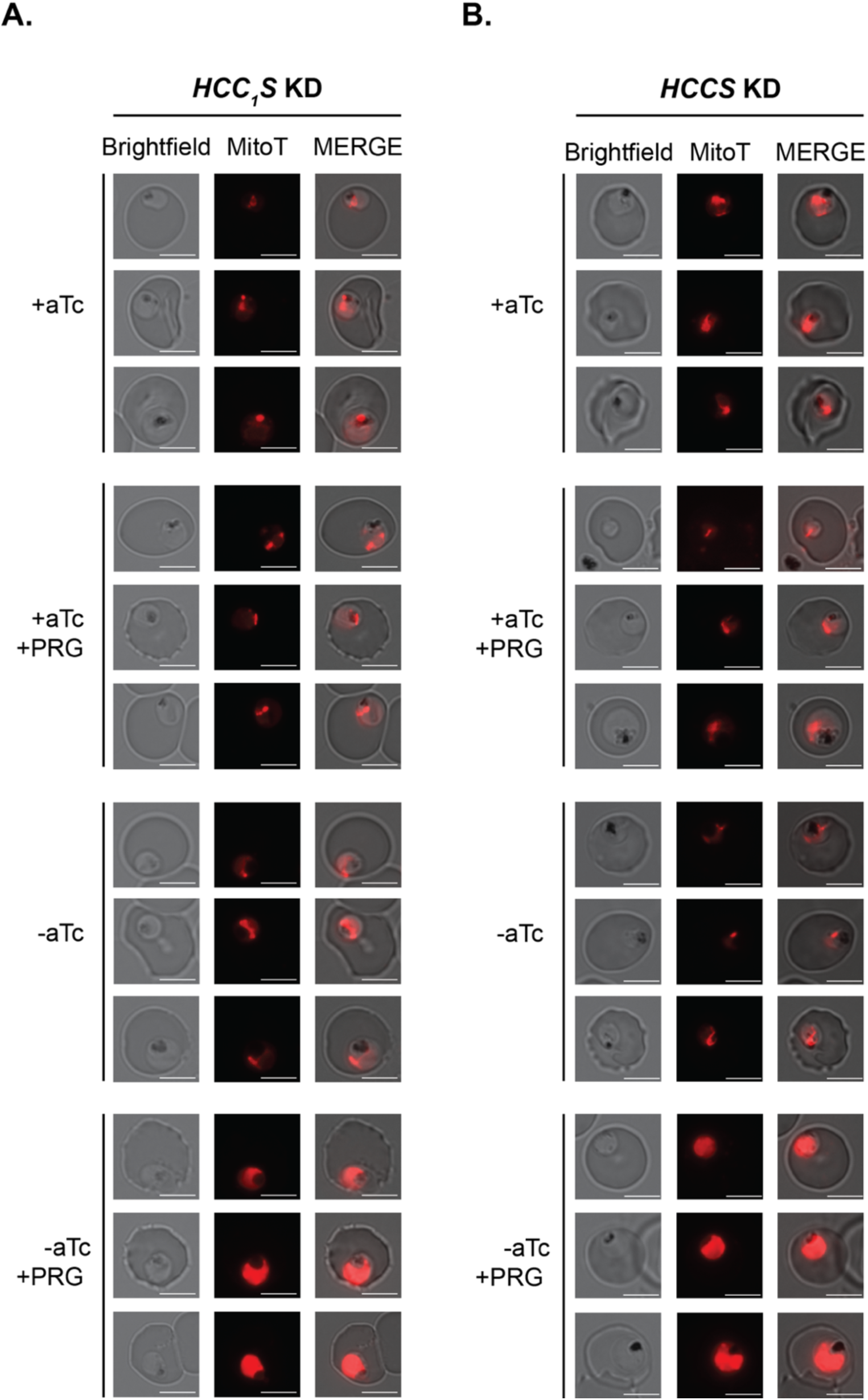
Additional live-parasite microscopy images of HCC_1_S (A) and HCCS (B) aptamer/TetR-DOZI Dd2 parasites cultured 5 days ±aTc/+dQ and 1 µM proguanil (PRG) and stained with 1 nM with MitoTracker Red (MitoT). Scale bars represent 5 µm.

**Figure S6.**
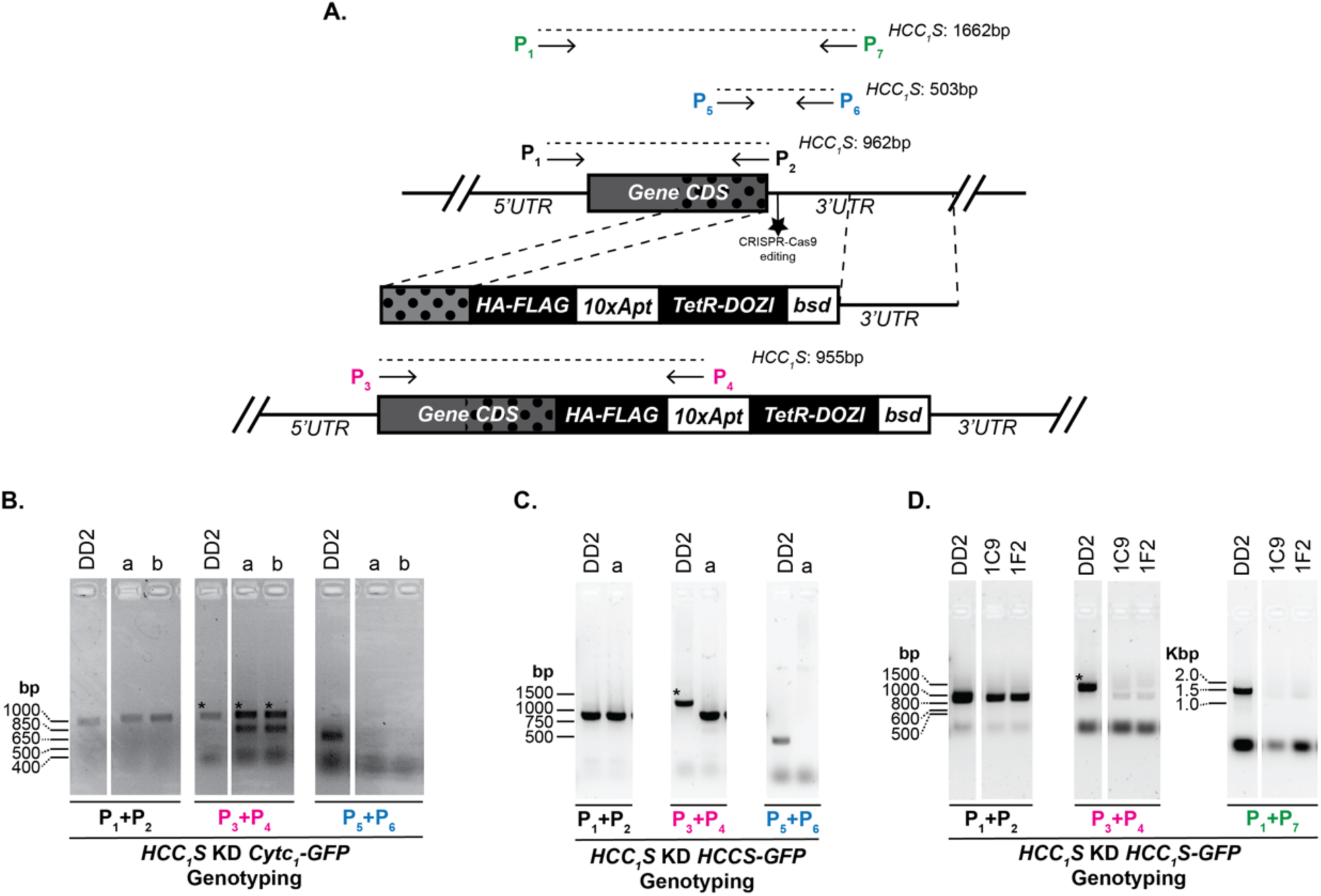
Genotyping HCC_1_S integrations in Dd2 parasites transfected with plasmids for episomal expression. (A) Schematic model for editing the chromosomal gene for *P. falciparum* HCC_1_S to encode a C-terminal HA-FLAG epitope tag fusion and the 10x aptamer/TetR-DOZI system. PCR primer sites are indicated with a “P” along with expected amplicon sizes for primer pairs. (B) Genotype PCR for parental and polyclonal transfectants (a and b) to edit HCC_1_S in parasites transfected with a plasmid for episomal expression of cyt *c*_1_-GFP. (C) Genotype PCR for parental and polyclonal transfectant a to edit HCC_1_S in parasites transfected with a plasmid for episomal expression of HCCS-GFP. (D) Genotype PCR for parental and clonal transfectants 1C9 and 1F2 to edit HCC_1_S in parasites transfected with a plasmid for episomal expression of HCC_1_S-GFP. Non-specific amplicons are indicated with asterisks.

**Figure S7.**
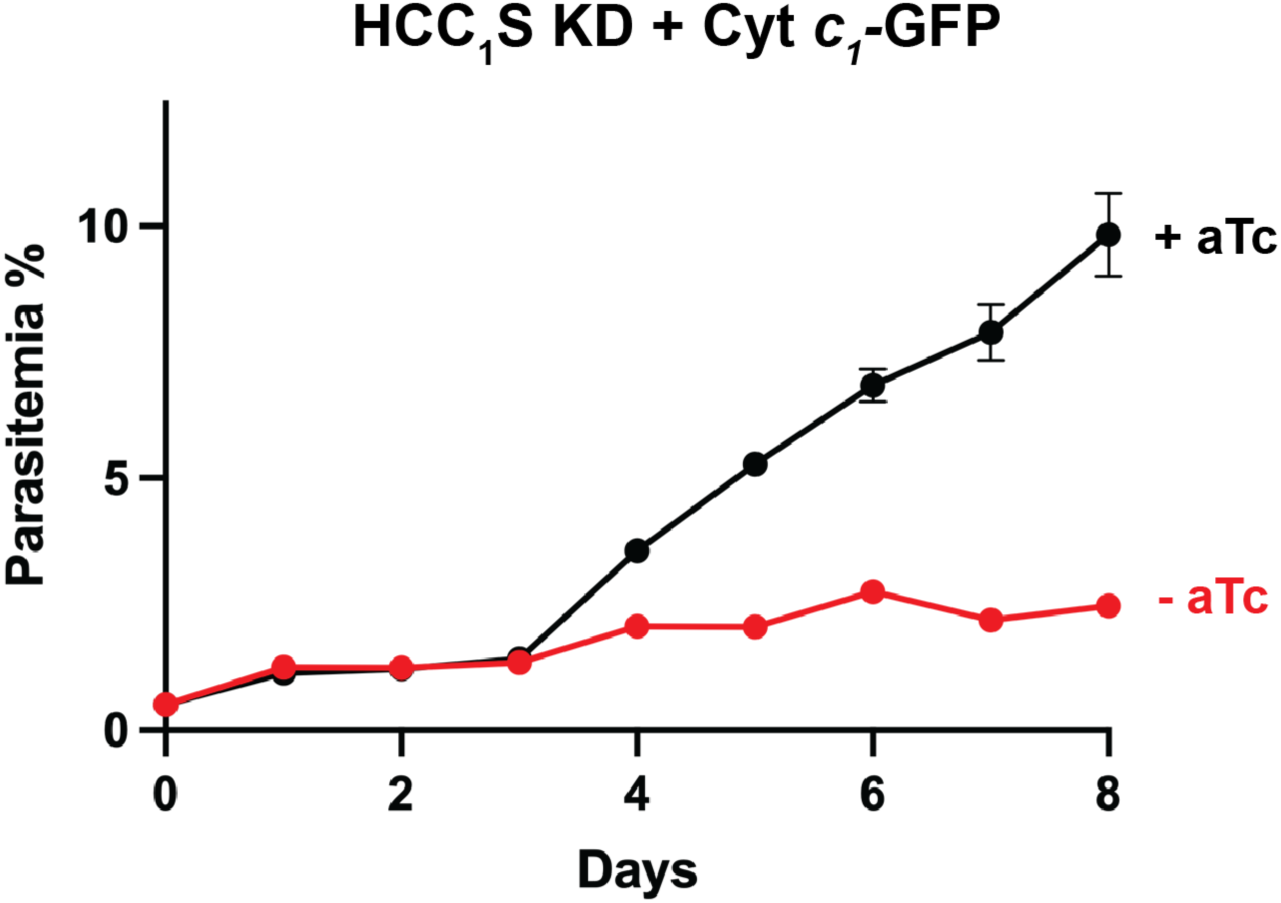
Synchronous growth assay of Dd2 parasites tagged for conditional knockdown (KD) of HCC_1_S expression in ±anhydrotetracycline (aTc) conditions and transfected with a plasmid for episomal expression of cyt *c*_1_-GFP. Data points and error bars represent the average ± SD of biological triplicates.

**Figure S8.**
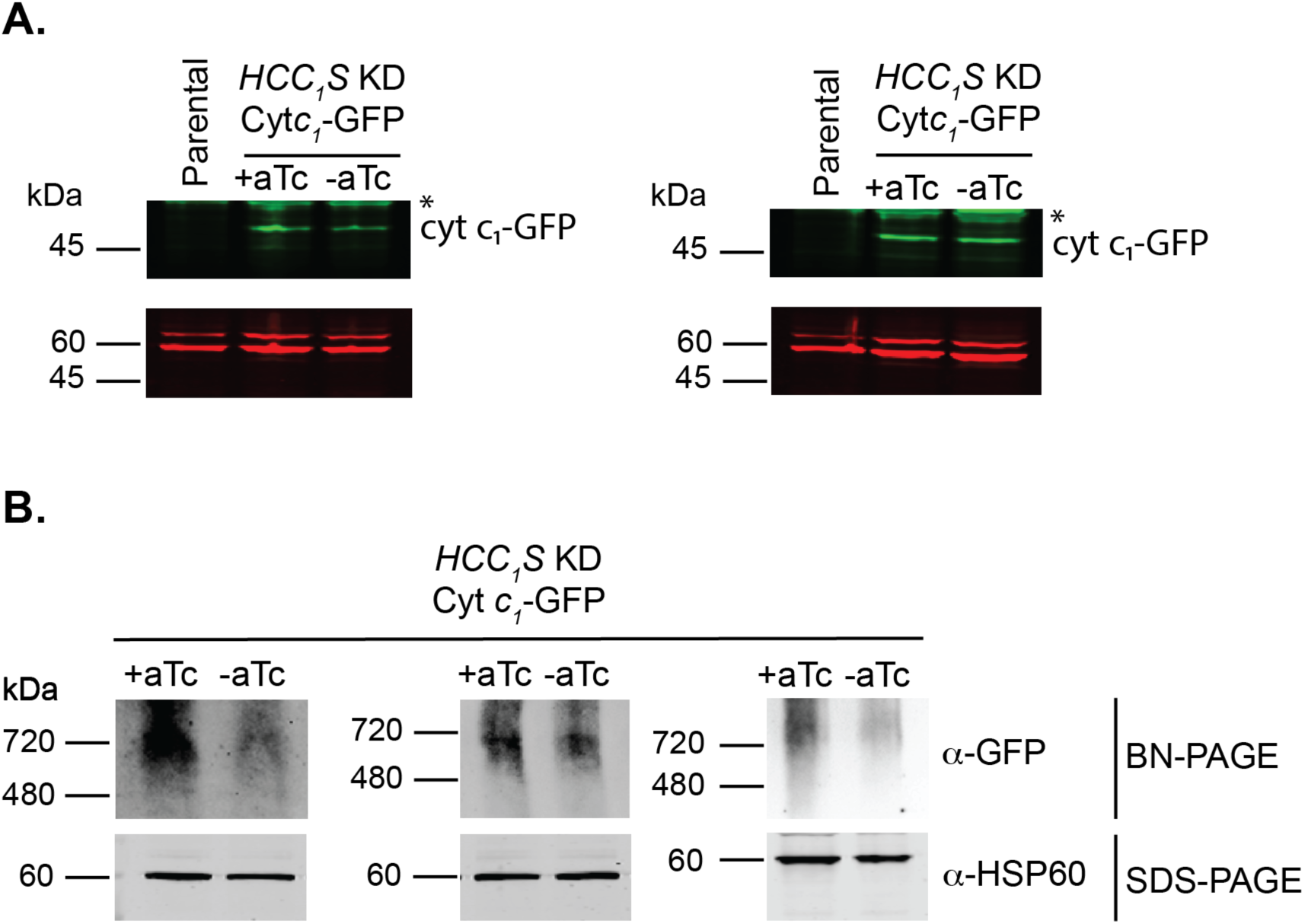
Additional western blot (WB) images from biological replicate growth assays of HCC_1_S knockdown Dd2 parasites episomally expressing cyt *c*_1_-GFP and grown 3 days ±aTc . (A) SDS-PAGE/WB images from independent growth assays (left and right) probed with α-GFP (green) and α-Hsp60 (red) antibodies. The asterisk indicates non-specific background bands visible in parental parasites (see Fig. S9 for uncropped images). (B) Blue native-PAGE/WB analysis of cyt *c*_1_-GFP and SDS-PAGE/WB analysis of HSP60 expression levels in HCC_1_S knockdown parasites cultured 5 days ±aTc/+dQ. 20 µg of total protein from mitochondrial lysates were loaded for each lane. Blots were probed with α-GFP or α-HSP60 antibodies.

**Figure S9.**
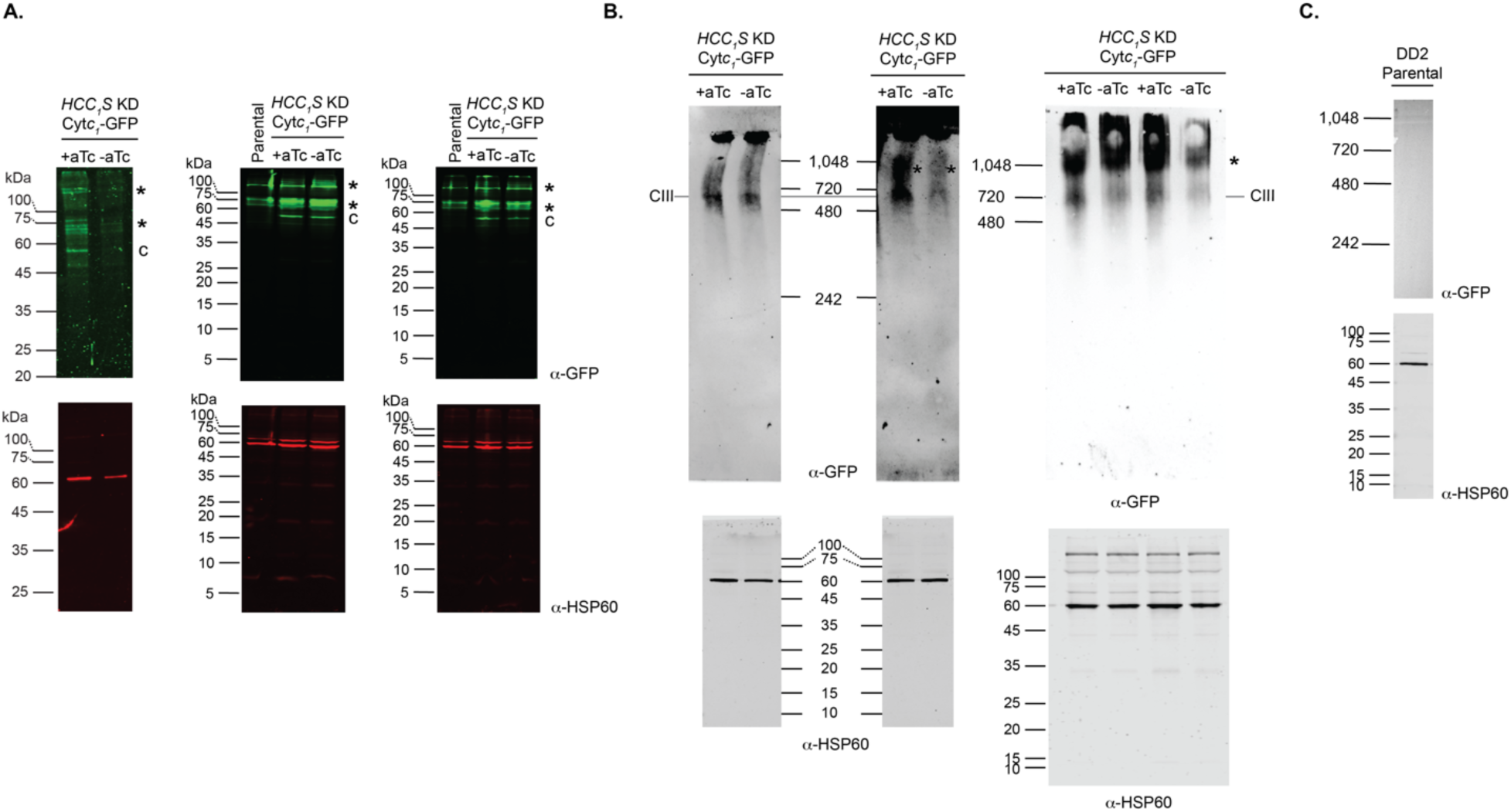
Uncropped western blot images. (A) SDS-PAGE/WB analysis of cyt *c*_1_-GFP levels in HCC_1_S knockdown (KD) Dd2 parasites cultured in ±aTc/+dQ conditions and probed with anti-GFP or anti-HSP60 antibodies. Asterisks label non-specific anti-GFP signal detected in parental parasites. The letter “c” indicates the specific band for cyt *c*_1_-GFP. (B) BN-PAGE/WB analyses of cyt *c*_1_-GFP levels in HCC_1_S knockdown (KD) Dd2 parasites cultured in ±aTc/+dQ conditions and probed with anti-GFP or anti-HSP60 antibodies. The lower panels are SDS-PAGE/WB analyses for HSP60 of identical samples as loaded in equivalent lanes in the upper panel images. Asterisks in reflect anti-GFP signal that does not correspond to the known size for Complex III ∼700 kDa. (C) The upper panel is a BN-PAGE/WB analysis of parental Dd2 parasites probed with an anti-GFP antibody. The lower panel is an SDS-PAGE/WB of an identical sample from parental Dd2 parasites probed with anti-HSP60 antibody.

**Figure S10.**
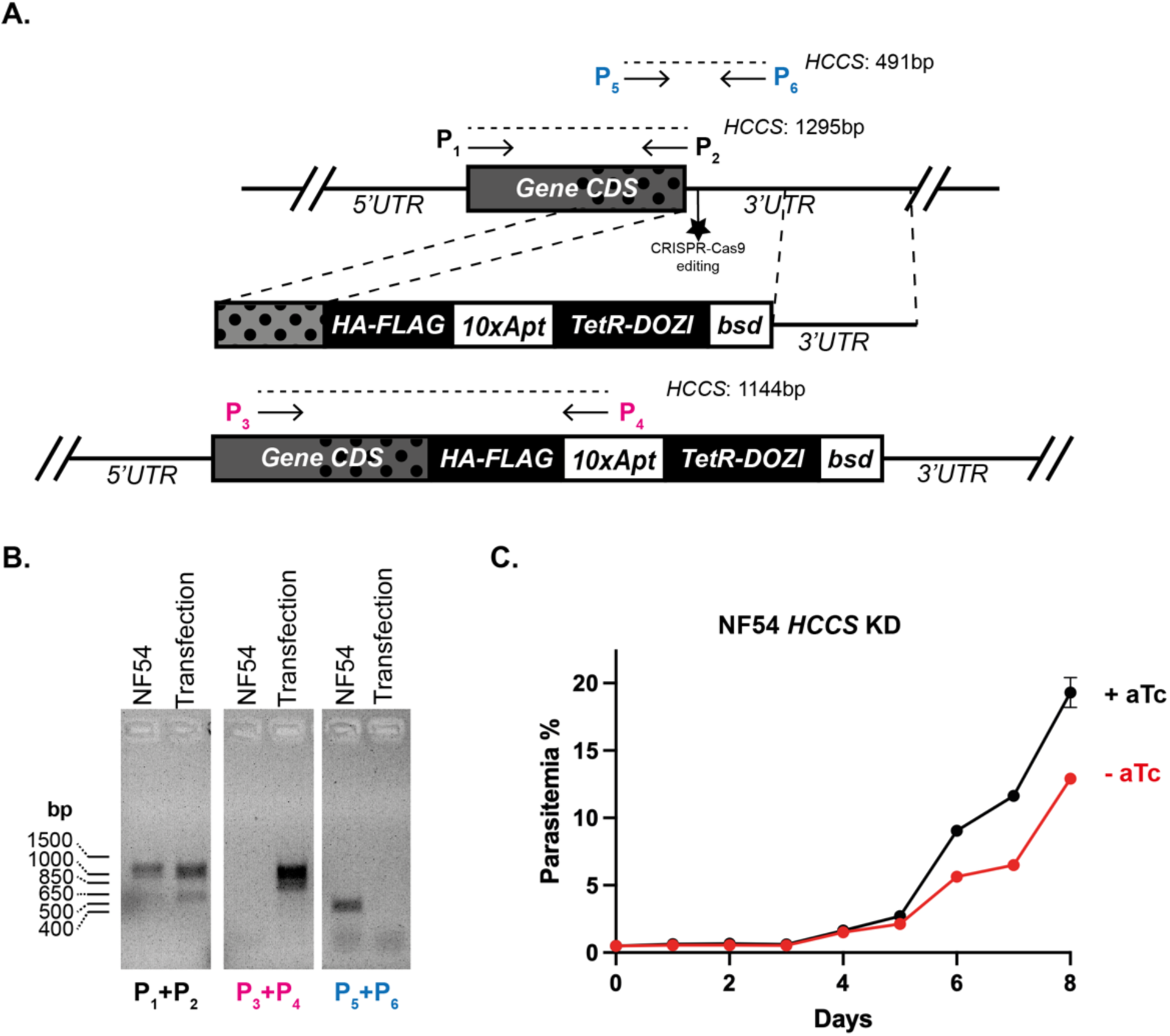
Genotyping HCCS integration in NF54 parasites. (A) Schematic model for editing the chromosomal gene for *P. falciparum* HCCS to encode a C-terminal HA-FLAG epitope tag fusion and the aptamer/TetR-DOZI system. PCR primer sites are indicated with a “P” along with expected amplicon sizes for primer pairs. (B) Genotype PCR for parental and polyclonal transfectants to edit HCC_1_S. (C) Synchronous growth assay of NF54 parasites tagged for conditional knockdown (KD) of HCCS expression in ±anhydrotetracycline (aTc) conditions. Data points and error bars represent the average ± SD of biological triplicates.

**Figure S11.**
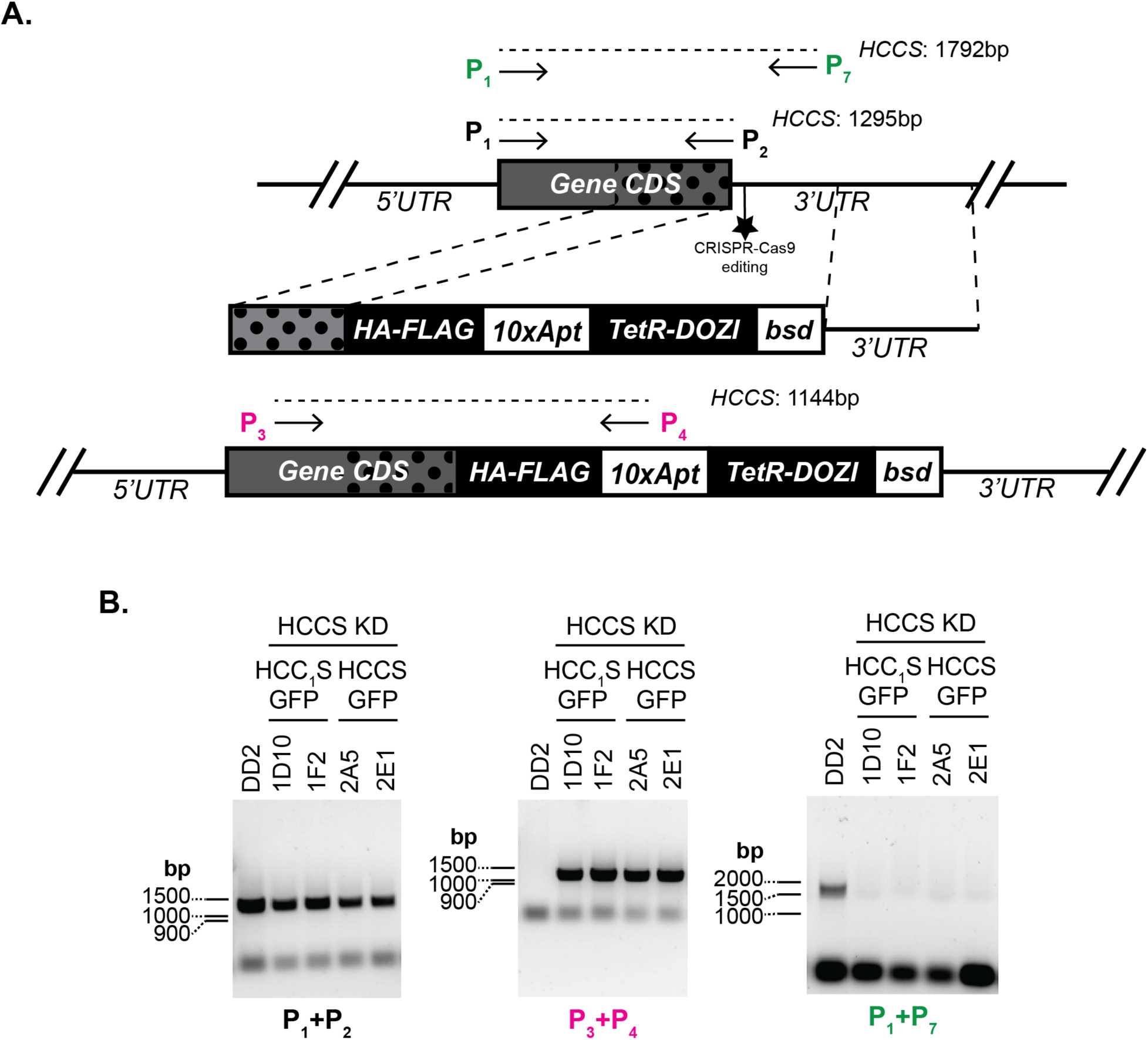
Genotyping HCCS integrations in Dd2 parasites transfected with plasmids for episomal expression. (A) Schematic model for editing the chromosomal gene for *P. falciparum* HCCS to encode a C-terminal HA-FLAG epitope tag fusion and the 10x aptamer/TetR-DOZI system. PCR primer sites are indicated with a “P” along with expected amplicon sizes for primer pairs. (B) Genotype PCR for parental and clonal transfectants to edit HCCS in parasites episomal expressing HCC_1_S-GFP or HCCS-GFP.

**Figure S12.**
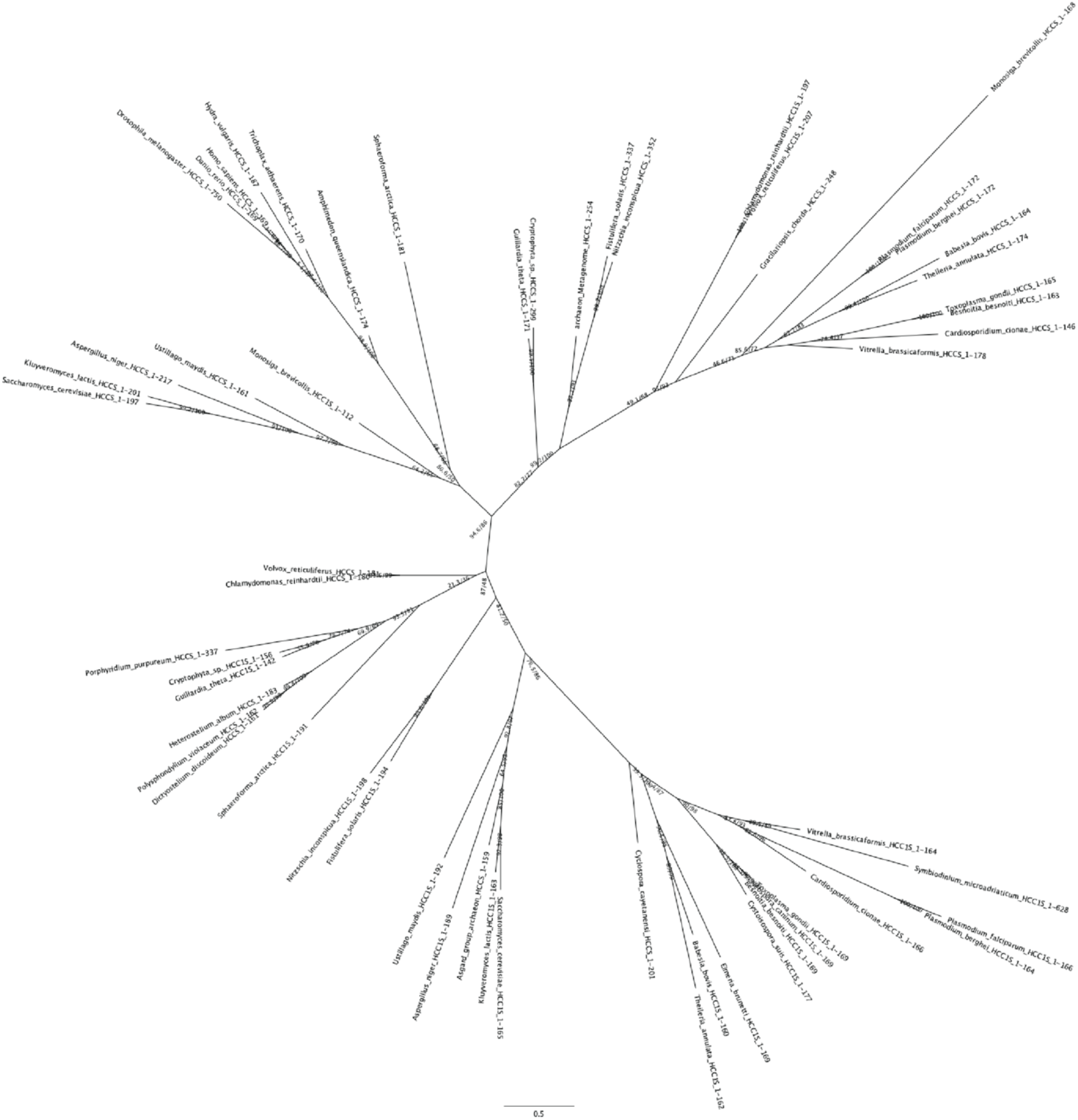
Raw phylogenetic tree based on sequence comparison of *P. falciparum* HCCS and HCC_1_S relative to diverse eukaryotes. This raw tree was used to create Figure 2B.

**Table S1.** List of organisms and HCCS/HCC_1_S protein sequences aligned in Fig. S12.

| Organism | GenBank |
| --- | --- |
| *Plasmodium falciparum* | KAF4327347.1 |
| *Plasmodium falciparum* | KAF4327070.1 |
| *Plasmodium berghei* | XP_034424219.1 |
| *Plasmodium berghei* | XP_034420526.1 |
| *Toxoplasma gondii* | KAF4646083.1 |
| *Toxoplasma gondii* | XP_002364614.2 |
| *Eimeria brunetti* | CDJ51885.1 |
| *Neospora caninum* | XP_003885328.1 |
| *Cystoisospora suis* | PHJ20471.1 |
| *Symbiodinium microadriaticum* | OLQ04208.1 |
| *Cyclospora cayetanensis* | XP_022588510.2 |
| *Babesia bovis* | XP_001612270.1 |
| *Babesia bovis* | EDO07243.2 |
| *Theileria annulata* | XP_951931.1 |
| *Theileria annulata* | XP_953889.1 |
| *Cardiosporidium cionae* | KAF8819734.1 |
| *Cardiosporidium cionae* | KAF8822908.1 |
| *Besnoitia besnoiti* | XP_029222774.1 |
| *Besnoitia besnoiti* | XP_029218356 |
| *Vitrella brassicaformis* | CEM02166.1 |
| *Vitrella brassicaformis* | CEL92815.1 |
| *Chlamydomonas reinhardtii* | XP_042915786.1 |
| *Chlamydomonas reinhardtii* | XP_042918551.1 |
| *Volvox reticuliferus* | GIL82651.1 |
| *Volvox reticuliferus* | GIL70918.1 |
| *Porphyridium purpureum* | KAA8493688.1 |
| *Gracilariopsis chorda* | PXF40648.1 |
| *Asgard group archaeon* | MCP8720244.1 |
| *archaeon (Metagenome)* | RYH06097.1 |
| *Dictyostelium discoideum* | XP_643563.1 |
| *Polysphondylium violaceum* | KAF2072366.1 |
| *Heterostelium album* | XP_020432267.1 |
| *Heterostelium album* | XP_020427510.1 |
| *Fistulifera solaris* | GAX15878.1 |
| *Fistulifera solaris* | GAX20154.1 |
| *Nitzschia inconspicua* | KAG7370135.1 |
| *Nitzschia inconspicua* | KAG7361072.1 |
| *Guillardia theta* | XP_005830278.1 |
| *Guillardia theta* | XP_005839063.1 |
| *Cryptophyta sp.* | KAJ1487503.1 |
| *Cryptophyta sp.* | KAJ1494728.1 |
| *Kluyveromyces lactis* | CAH01636.1 |
| *Kluyveromyces lactis* | CAG98941.1 |
| *Ustilago maydis* | XP_757596.1 |
| *Ustilago maydis* | XP_760849.1 |
| *Aspergillus niger* | XP_001400653.1 |
| *Aspergillus niger* | XP_001401520.1 |
| *Saccharomyces cerevisiae* | NP_009361.1 |
| *Saccharomyces cerevisiae* | NP_012836.1 |
| *Drosophila melanogaster* | AAF55946.1 |
| *Danio rerio* | NP_958859.1 |
| *Homo sapiens* | NP_001116080.1 |
| *Monosiga brevicollis* | XP_001749606.1 |
| *Monosiga brevicollis* | XP_001748514.1 |
| *Sphaeroforma arctica* | XP_014157735.1 |
| *Amphimedon queenslandica* | XP_003385371.1 |
| *Trichoplax adhaerens* | XP_002114862.1 |
| *Hydra vulgaris* | XP_047122850.1 |

**Table S2.**
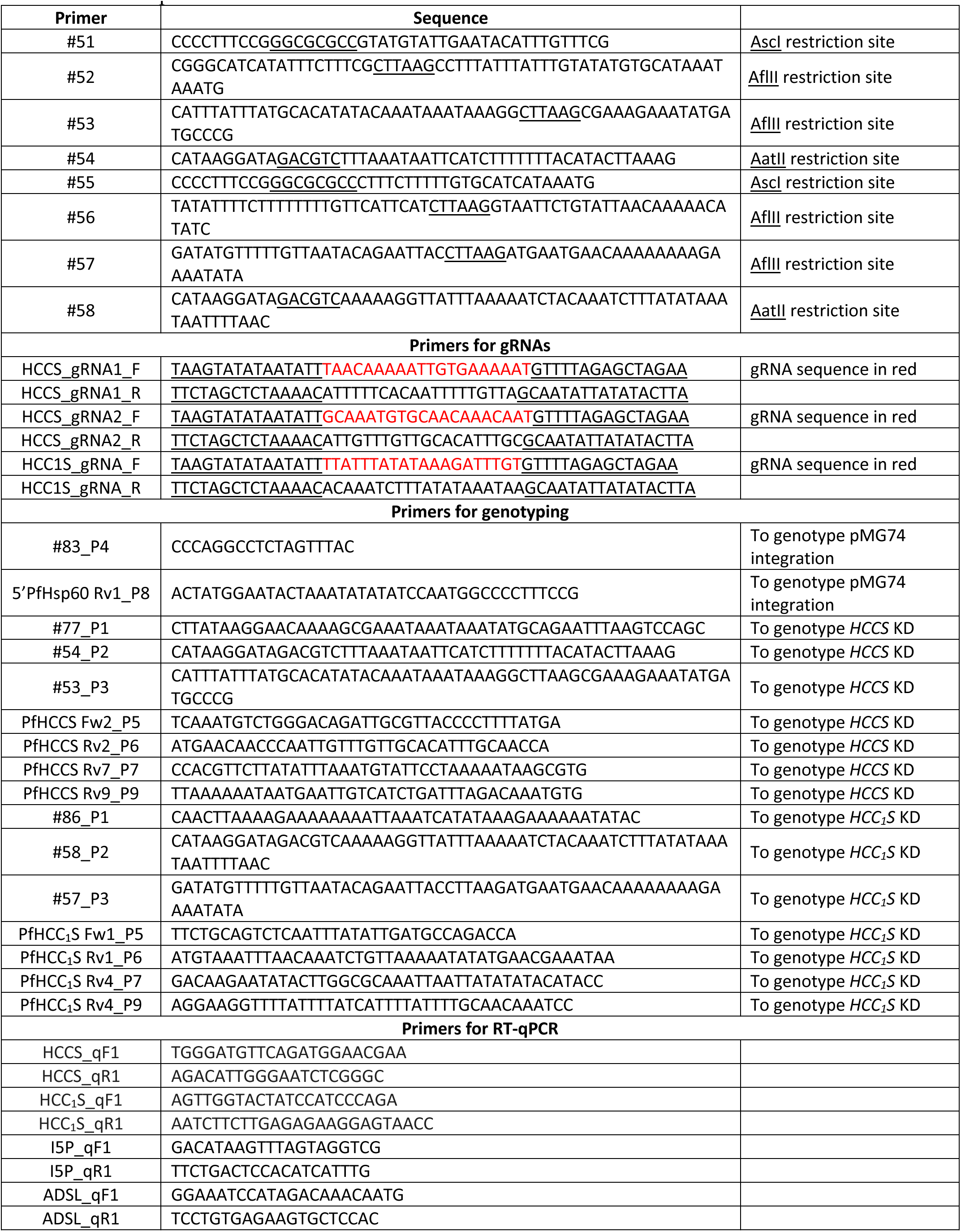
List of primers.

